# Coincidence detection in aspiny interneuron dendrites

**DOI:** 10.1101/2025.09.10.675107

**Authors:** Simonas Griesius, Amy Richardson, Marion Mercier, Dimitri M Kullmann

## Abstract

Inhibitory interneurons are conventionally thought to support precise timing of neural activity and regulation of circuit excitability, whilst principal neurons are the main locus of synaptic plasticity. Recent evidence for long-term potentiation (LTP) at glutamatergic synapses on GABAergic interneurons calls for close attention to the sub-cellular phenomena that underlie the induction of plasticity in aspiny dendrites. Neurogliaform interneurons, in particular, exhibit large NMDA receptor-mediated currents, robust Hebbian LTP and pronounced supralinear summation of glutamatergic signals converging on dendritic compartments. Unlike pyramidal neurons, however, dendritic supralinearity in interneurons operates independently of sodium channels. Here we investigate the principles governing supralinear synaptic integration and how it interacts with backpropagating action potentials in the context of synaptic plasticity.

We developed a biophysically realistic, multi-compartmental model of a murine hippocampal neurogliaform interneuron, with dendritic architecture and ion channel densities tuned to match *ex vivo* recordings. The interaction of synaptic inputs and action potential were validated with patch-clamp electrophysiology and two-photon imaging of dendritic calcium-dependent fluorescence transients.

We show that, whether simulated glutamatergic excitatory postsynaptic potentials are clustered on a dendritic fragment or dispersed across the entire arbor, they summate supralinearly as assessed from the calculated somatic voltage response. Clustered inputs however induce more pronounced local depolarizations and calcium transients than dispersed inputs. Action potentials generated at the soma backpropagate throughout the dendritic arbor but are attenuated at branch points. Coincident synaptic input and backpropagating action potentials relieve this attenuation, further enhance dendritic depolarizations, and greatly potentiate calcium influx via NMDA receptors. Despite differences in dendritic anatomy and in the roles of sodium channels, the interplay of orthodromic and antidromic voltage propagation and resultant calcium signal amplification exhibits remarkable similarities between neurogliaform interneurons and principal cells.

**Author summary:** Learning and memory are believed to rely in large part on the brain’s ability to strengthen excitatory connections among principal cells. Potentiation of excitation must however be balanced by inhibition. Our study focuses on a key player in this balancing act: the neurogliaform interneuron, which exerts a powerful inhibition of the apical dendrites of principal neurons in the cortex and hippocampus. Recent work has revealed that both principal neurons and neurogliaform interneurons amplify coincident excitatory signals in their extensively branching dendrites. Key to the induction of synaptic potentiation is the coincidence of orthodromic depolarization arising from synapses and action potentials generated at or near the cell body. We asked how these electrical signals interact, using a blend of computer simulations and experimental manipulations. Unlike principal neurons, neurogliaform interneurons are devoid of dendritic spines, and voltage-gated sodium channels are dispensable for boosting of synaptic depolarization. Nevertheless, the core principle that coincident orthodromic and antidromic signals leads to widespread depolarization spreading throughout the dendritic arbor is conserved between these neurons. The conclusions of this work shed light not only on healthy brain function but also on common disorders such as epilepsy and schizophrenia, which abnormal dendritic function has been implicated.

## Introduction

Dendrites receive incoming synaptic inputs, integrate them, and transmit information toward the soma (1). Neuronal signals can also travel through the dendrites in the opposite direction in the form of backpropagating action potentials (2–4). These bidirectional dendritic depolarizations may also interact with each other (5–8), thereby increasing neuronal computational capacity, including the ability to act as multi-level networks (9–14), to amplify signals (15), detect coincidence (16), perform XOR logic gating (17, 18), and compute prediction errors (13, 19).

Another important consequence of the interplay of forward and backward signalling in dendrites is that it opens a window for the induction of long-term potentiation (LTP) of synaptic strength. This is conventionally explained by NMDA receptors (NMDARs) acting as coincidence detectors because their opening and consequent calcium influx requires both presynaptic glutamate release and postsynaptic depolarization (16, 20, 21).

Although dendritic nonlinearities and NMDAR-dependent LTP have been studied extensively in pyramidal neurons, recent studies have revealed that GABAergic interneurons also perform complex dendritic operations (22–29), and robust NMDAR-dependent LTP can be elicited in some subtypes of GABAergic cells (30–32).

Despite the similarities between NMDAR-dependent LTP in interneurons and pyramidal neurons, their dendrites exhibit some striking anatomical and functional differences. Interneuron dendrites are typically aspiny or only have sparse ‘stubby’ spines, unlike pyramidal neurons (33). Moreover, supralinear summation of glutamatergic inputs to interneuron dendrites does not involve voltage-gated sodium channels (22–24), which in contrast contribute substantially to dendritic spikes and back-propagation of action potentials in pyramidal neurons (34–36).

Given the importance of interneurons to brain circuit and network excitability, rhythmicity, computations, and modulation (37–41), and likely involvement of active dendrites (22–29), we undertook the present study, based on computer simulations, electrophysiology and calcium imaging, to provide an understanding of non-linear voltage signalling that cannot be obtained from patch-clamp recordings alone. We focus on neurogliaform cells, which contribute to feed-forward inhibition of apical tufts of principal cells (42, 43) and exhibit excitatory postsynaptic potentials and currents (EPSPs and EPSCs) with an especially large NMDA:AMPA ratio (42). Neurogliaform cells also exhibit robust NMDAR-dependent forms of long-term potentiation (LTP) (30). We recently reported that neurogliaform interneurons summate spatially clustered near-synchronous dendritic inputs supralinearly (22). This phenotype was identified to be critically dependent on NMDARs, with L-type voltage-gated calcium channels (VGCCs), and intracellular calcium stores contributing downstream to boost calcium signals in dendrites (22).

To investigate how synaptic inputs interact with backpropagating action potentials in the context of different forms of synaptic plasticity, we performed biophysically detailed simulations in a reconstructed murine neurogliaform interneuron. We reveal some similarities between the principles underlying the interplay of synaptic depolarization and backpropagating action potentials, including the ability to boost NMDAR-mediated calcium influx despite the absence of involvement of sodium channels, but also some subtle differences in the ability to compartmentalize such signals. To complement our computational modelling and directly link dendritic integration to synaptic plasticity, we additionally performed a secondary analysis of previously published experimental data (30). In that study, theta-burst stimulation (TBS) of the temporoammonic pathway produced robust LTP in neurogliaform interneurons, even in the absence of imposed somatic depolarization (30). This analysis was motivated by the hypothesis that variability in LTP induction may reflect differences in dendritic integration of synaptic inputs, rather than simply differences in baseline synaptic strength. We also test our *in silico* predictions using new 2-photon calcium imaging *ex vivo* experiments on backpropagating action potentials. We show that neurogliaform interneurons readily integrate temporally coherent converging synaptic inputs, whether clustered or dispersed, and perform non-linear computations on forward- and backpropagating voltage and calcium signals

## Results

### The multi-compartment model recreates the key experimental phenotype of supralinear dendritic integration

We first sought to create an *in silico* model that captured the key dendritic integration phenotypes of neurogliaform interneurons reported experimentally (22, 30, 42). The model was based on a morphological reconstruction of the same neurogliaform interneuron previously characterised experimentally in Griesius et al. (22), from which electrophysiological and calcium imaging data were also obtained. The example neuron was reconstructed and segmented into compartments (**Figure 1a**, **Supplementary Figure S1**) and imported into the NEURON simulation environment (44). Passive and active parameters were set based on values taken from other studies on dendritic integration (14, 22, 30, 42, 45–50) (**Tables 1-3**, see Methods for details).

**Figure 1.**
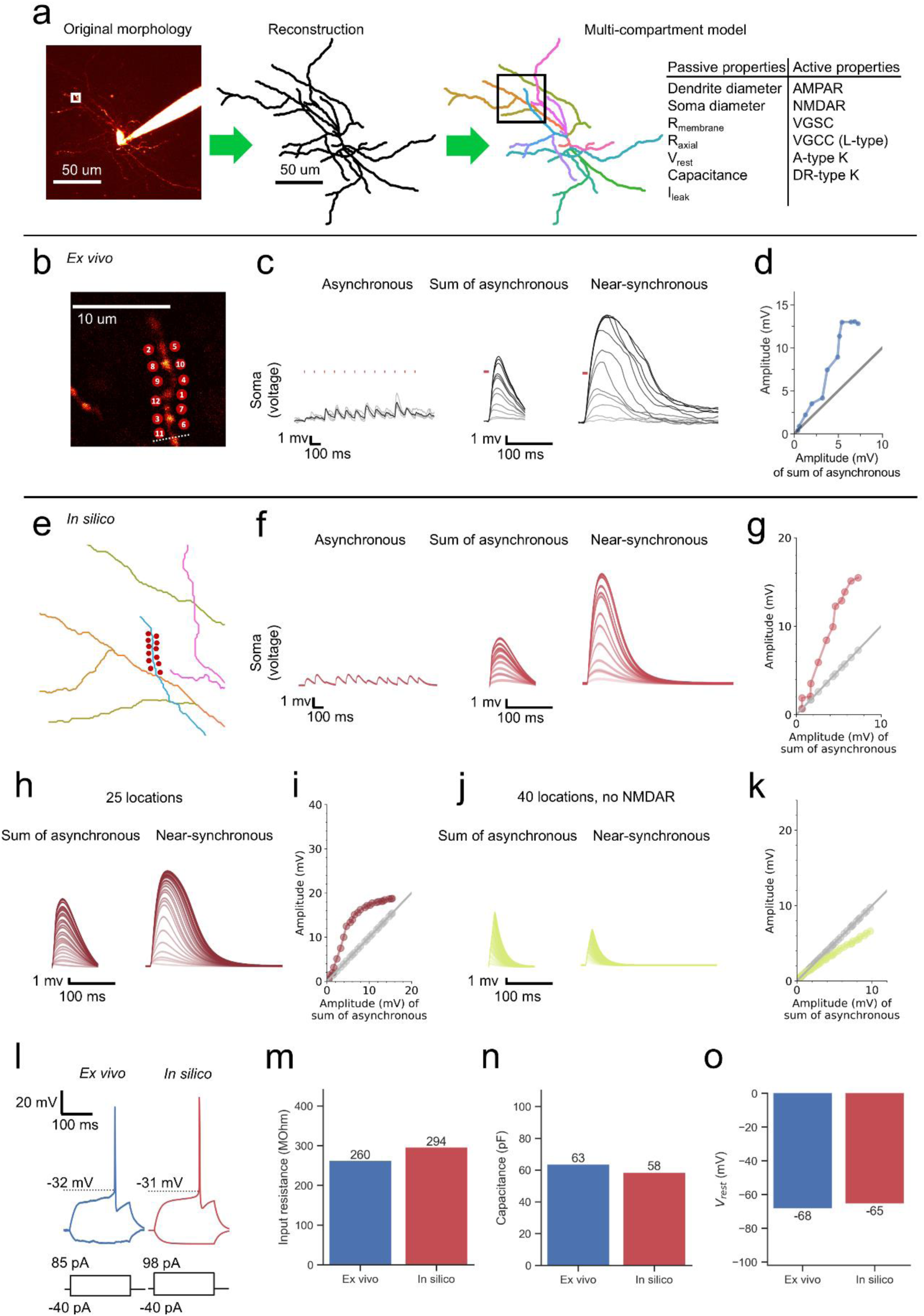
The multi-compartment *in silico* model recreates the key *ex vivo* experimental phenotype of NMDAR-dependent supralinear dendritic integration. **(a)** Model overview. A representative neurogliaform interneuron patch-filled with the inert dye Alexa-594. It was reconstructed and segmented into a multi-compartment model populated with passive and active properties. **(b)** Inset showing locations used for 2-photon glutamate uncaging *ex vivo*. The inset corresponds to the location shown in (a). **(c)** Somatic voltage uEPSP traces across asynchronous and near-synchronous conditions. The sum of asynchronous uEPSPs is also displayed. Laser uncaging light stimuli are indicated by red dots above the traces. uEPSPs were elicited in the asynchronous condition with light pulses separated by an interval of 100.32ms (left; grey traces: individual sweeps; black traces average). The sum of asynchronous uEPSP traces were calculated by aligning and summing an increasing number of asynchronously evoked uEPSPs. Near-synchronous uEPSPs were elicited with light pulses separated by an interval of 0.32ms (right). The families of traces for the sum of asynchronous and near-synchronous uEPSPs are shown as light to dark grey traces, indicating an increase in the number of uncaging loci from 1 to 12. **(d)** Amplitude of the recorded near-synchronous uEPSP plotted against the amplitude of the sum of asynchronous uEPSPs as the number of uncaging loci was increased. The line of identity is represented by a grey line. Data from Griesius et al. 2025, reused with author permission (22). **(e)** Inset showing locations used for simulating synaptic input in the *in silico* multi-compartment model. The inset corresponds to the location shown in (a). **(f)** Simulated somatic voltage traces across asynchronous and near-synchronous conditions. Displayed as in (c). **(g)** Amplitude of the simulated near-synchronous uEPSP plotted against the amplitude of the sum of asynchronous uEPSPs as the number of activated synapses was increased. The line of identity is represented by a grey line. **(h)** The same simulation as in (f) but with a total of 25 synaptic locations activated. **(i)** Amplitude of the simulated near-synchronous uEPSP plotted against the amplitude of the sum of asynchronous uEPSPs as the number of activated synapses was increased. The line of identity is represented by a grey line. **(j)** The same simulation as in (h) but with NMDARs removed. **(k)** Amplitude of the simulated near-synchronous uEPSPs plotted against the amplitude of the sum of asynchronous uEPSPs as the number of activated synapses was increased. The line of identity is represented by a grey line. **(l)** Rheobase recorded from the representative neurogliaform interneuron *ex vivo* and from the *in silico* model derived from it. Action potential thresholds are indicated by dashed lines. Current injection steps and their amplitudes are shown below the corresponding voltage traces. **(m-o)** Comparison of intrinsic membrane properties between the *ex vivo* neuron and the corresponding *in silico* model, including input resistance **(m)**, membrane capacitance **(n)**, and resting membrane potential **(o)**. Values are shown for the same neuron used for reconstruction.

**Table 1.** Comparison of intrinsic properties between *ex vivo* recording and its *in silico* model.

| Intrinsic property | *Ex vivo* | *In silico* |
| --- | --- | --- |
| **Ri (MΩ)** | 260 | 294 |
| **Rheobase (pA)** | 85 | 98 |
| **V<sub>rest</sub> (mV)** | -67.6 | -65.3 |
| **Spike threshold (mV)** | -32 | -31 |
| **Spike half-width (ms)** | 1.7 | 1.7 |
| **Capacitance (pF)** | 62.8 | 57.5 |
| **Tau (ms)** | 16.3 | 16.8 |

Ri Input resistance
V<sub>rest</sub> Resting membrane potential

In our *ex vivo* study (22) dendritic summation of synaptic signals was explored by 2-photon uncaging glutamate at 12 closely-spaced locations along a dendritic fragment either asynchronously (100.32 ms interval between locations) or near-synchronously (0.32 ms interval). Individual uncaging EPSPs (uEPSPs) evoked with asynchronous stimulation and recorded at the soma (**Figure 1b-c**) were summated and compared to the uEPSP in response to near-synchronous glutamate uncaging. The experimentally observed uEPSP evoked by near-synchronous uncaging was consistently greater than predicted by the sum of the separate asynchronously-evoked uEPSPs and was therefore supralinear (**Figure 1c-d**). We asked if this supralinear dendritic response can be recreated *in silico*.

The same dendrite that was stimulated experimentally using 2-photon glutamate uncaging was identified (dendrite 35) and populated *in silico* with 12 AMPA receptor (AMPAR) and NMDAR conductance-based point processes to simulate 12 closely spaced glutamatergic synapses (**Figure 1e**). When these synapses were activated asynchronously to produce separate responses, the resulting uEPSPs at the soma were of similar magnitude to their *ex vivo* counterparts (∼1-2 mV) (**Figure 1f**). When the same synapses were activated near-synchronously, the uEPSP at the soma summated supralinearly (**Figure 1f-g**), recreating the key experimental phenotype, suggesting the model could be used to make additional biophysically plausible predictions on neurogliaform dendritic computation.

The inclusion of 13 additional synapses onto the same dendrite 35 (25 synapses total) revealed saturation of supralinearity (**Figure 1h-i**). The uEPSP amplitude converged toward the linear prediction, generated from the sum of asynchronous uEPSPs, as expected from a collapse of the driving force under these conditions. Further, when the same simulation was repeated without NMDARs present, the near-synchronous uEPSPs at the soma never exhibited supralinearity (**Figure 1j-k**). The uEPSPs instead rapidly became sublinear, likely due to the collapse of the local driving force and shunting via open AMPARs. In line with experimental findings (22), the *in silico* model was tuned to have a relatively small contribution of voltage-gated sodium channels (VGSCs) and voltage-gated calcium channels (VGCCs) (**Supplementary Figure S2**, see Methods for details).

To validate the physiological fidelity of the model, intrinsic membrane properties were directly compared between the *in silico* reconstruction and *ex vivo* recordings obtained from the same neuron (22). Input resistance, resting membrane potential, membrane capacitance, and membrane time constant, rheobase, spike threshold, and spike half-width were all closely matched (**Figure 1l-o**; **Table 1**), demonstrating that the *in silico* model reproduces both passive membrane properties and excitability of the recorded cell. This agreement supports the use of the model for the mechanistic investigation of dendritic integration.

To assess the robustness of supralinear dendritic integration to changes in synaptic strength, we repeated the asynchronous and near-synchronous synaptic input simulations on dendrite 35 but this time with synaptic weights uniformly scaled down or up (0.5x and 2x, respectively) (**Supplementary figure S3**). In both cases, the magnitude of uEPSP supralinearity during near-synchronous stimulation was reduced compared to baseline conditions. When synaptic weights were decreased (0.5x), synaptic depolarization was insufficient to strongly recruit voltage-dependent conductances, resulting in a more linear summation of uEPSPs. Conversely, with larger synaptic weights (2x), the reduced driving force led to a collapse of the nonlinear amplification observed under baseline conditions. Despite these changes in amplitude, the overall pattern of supralinear integration during near-synchronous input was qualitatively preserved, albeit attenuated.

We further tested whether the supralinear dendritic integration readout observed in the *in silico* model depended on the specific reconstructed morphology by repeating the key simulations in two additional neurogliaform interneuron reconstructions (**Supplementary Figures S5–S6**). These additional reconstructions were also constrained by *ex vivo* physiology data recorded in these very neurons. Synaptic weights were adjusted to reproduce comparable asynchronous uEPSP amplitudes. In both cases, near-synchronous synaptic activation produced supralinear summation of uEPSPs, consistent with the primary model. While the magnitude of nonlinearity varied between morphologies, the effect was qualitatively unchanged. These results indicate that supralinear integration is a robust feature across neurogliaform interneuron morphologies rather than an artefact dependent upon fine-tuning a single reconstruction.

### Spatially clustered input onto a single dendrite enables robust supralinear summation

Although active synapses can be clustered *in vivo* (51, 52), they may also be dispersed across dendritic arbors (53, 54). It is challenging to systematically test the spatial arrangements of synaptic inputs *in vivo*, especially in deeper structures like the hippocampus. It is also technically difficult to compare widely different spatial configurations of synaptic inputs, be it *in vivo* or *ex vivo*. An advantage of *in silico* modelling is the ability to simulate and scrutinise the spread of depolarization throughout the entire dendritic tree.

We first hypothesized that the dendritic voltage supralinearities may invade a broad section of the dendritic arbor due to the relatively compact size of neurogliaform interneuron dendritic arbors. Indeed, when the same 12 glutamatergic synapses were activated near-synchronously, the elicited uEPSP was maximal in the activated dendrite 35, and attenuated with distance, not only towards the soma but also throughout the dendritic tree (**Figure 2a-b**). The amplitude at dendrite 31, which shares a parent dendrite with dendrite 35, exhibited the second-largest amplitude, followed by the soma, and two other sampled dendrites (6 and 12) (**Figure 2b**). The fast initial peak seen in dendrite 35 was AMPAR-mediated (**Supplementary figure S4**) and particularly susceptible to dendritic filtering due to the brevity of the event. Despite the attenuation with distance, the degree of supralinear summation, estimated by comparing the response amplitude 20 ms after synaptic stimulation onset to the sum of individual asynchronous uEPSPs, was only slightly lower in the sampled remote dendrites (median: 125%) than in dendrite 35, which received the active synapses (144%) (**Figure 2g**).

**Figure 2.**
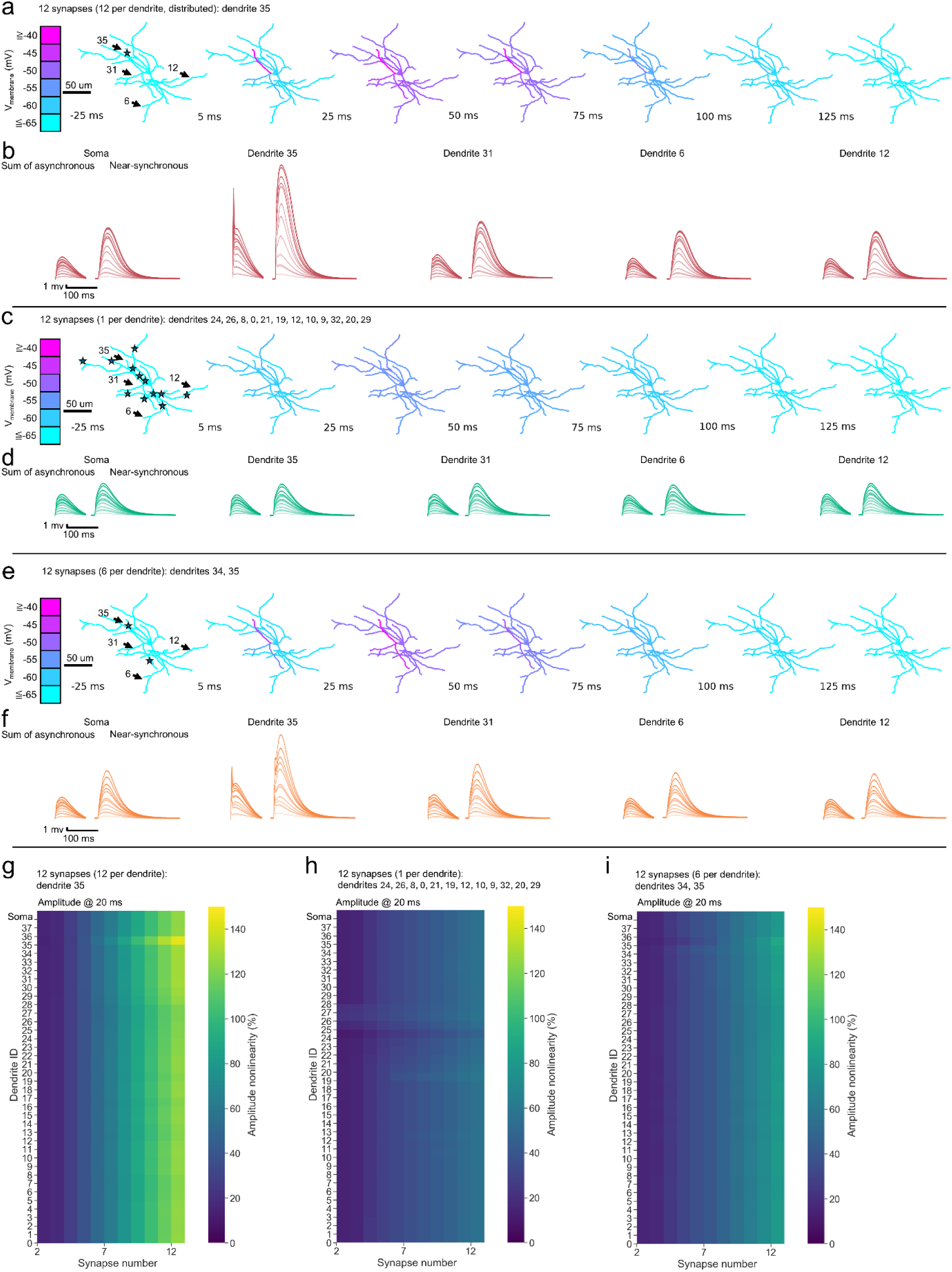
Spatially clustered input onto a single dendrite enables robust supralinear summation. **(a)** Simulated clustered synaptic input. Twelve synapses were activated on dendrite 35 (the same as in figure 1). Voltage snapshots at 25 ms before stimulation and 5-125 ms after stimulation onset overlaid onto the neuronal architecture. Stimulation site on dendrite 35 is indicated by the star. Dendrites 35, 31, 6, 12 are indicated with arrows and have their uEPSPs displayed as traces below. **(b)** Sum of the asynchronous and near-synchronous uEPSPs at the soma and dendrites 35, 31, 6, 12. **(c)** Simulated dispersed synaptic input. 12 synapses (1 per dendrite) were activated on dendrites 24, 26, 8, 0, 21, 19, 12, 10, 9, 32, 20, 29. Data displayed as in (b). **(d)** Sum of the asynchronous and near-synchronous uEPSPs at the soma and dendrites 35, 31, 6, 12. **(e)** Simulated clustered synaptic input on two nearby branches. 12 synapses (6 per dendrite) were activated on dendrites 35 and 34. Data displayed as in (b). **(f)** Sum of the asynchronous and near-synchronous uEPSPs at the soma and dendrites 35, 31, 6, 12. **(g)** Heatmap of amplitude nonlinearity for every dendrite and the soma. Corresponds to (a, b). **(h)** Heatmap of amplitude nonlinearity for every dendrite and the soma. Corresponds to (c, d). **(i)** Heatmap of amplitude nonlinearity for every dendrite and the soma. Corresponds to (e, f). Amplitude was measured at 20 ms post stimulus onset in (g-i).

Next, we hypothesized that due to their relatively small size, neurogliaform interneurons may be able to summate uEPSPs supralinearly in response to near-synchronous synaptic input even when the input is distributed across the dendritic arbor. Therefore, 12 glutamatergic synapses were distributed across a random subset of dendrites (1 synapse per dendrite) and the same synaptic stimulation timing was simulated (**Figure 2c).** The resulting depolarization again spread widely through the dendritic tree. Under these simulation conditions, however, the near-synchronous uEPSP amplitude was now substantially reduced compared to the uEPSP amplitude arising from the activation of synapses on a single dendrite (**Figure 2d**). Interestingly, the near-synchronous synaptic inputs summated supralinearly even with activated synapses spread across different dendrites (median supralinearity 55%), albeit to a lesser degree than when they were clustered (125%) (**Figure 2h**). The maximal supralinearity recorded under this paradigm emerged in dendrite 19 (60%), possibly due to dendritic morphology and the chance sequence of activated dendrites.

When 12 glutamatergic synapses were dispersed among 2 nearby dendrites (dendrites 34 and 35) and the same synaptic stimulation timing was simulated, the overall picture was somewhere in between the previous two simulated conditions (**Figure 2e, f, i**). Near-synchronous stimulation generated uEPSPs that were greater in amplitude than predicted by the sums of asynchronous uEPSPs. The near-synchronous uEPSP amplitude was the greatest in the activated dendrites, from where the depolarization spread throughout the dendritic tree and attenuated with distance. Likewise, the degree of supralinearity was the greatest in the two dendrites with the activated synapses (dendrite 34: 86%; dendrite 35: 90%) (**Figure 2i**). The median degree of supralinearity across dendrites (80%) fell between that seen when all activated synapses were concentrated on a single dendrite (125%) and that seen when the activated synapses were completely dispersed (55%).

### Spatially dispersed inputs can elicit dendritic supralinearities

The simulation with activated synapses distributed across 12 dendrites suggested that supralinear summation is attenuated but nevertheless possible even with spatially dispersed inputs (**Figure 2c-d**). As more synapses were distributed across dendrites, this phenomenon persisted (**Supplementary Figure S7a-b**). The uEPSP amplitude evoked by near-synchronous synapse activation was consistently greater than the sum of asynchronous uEPSPs. Upon sufficient near-synchronous depolarization (21 synapses activated) a single action potential was evoked, which spread throughout the dendritic tree along with the uEPSP. The median degree of supralinearity across all dendrites was 53% and the maximum was 60% at dendrite 12 (**Figure S7c**), again notably smaller than when fewer inputs were clustered on a single or two nearby dendrites.

### Dendritic integration may drive theta burst stimulation-induced LTP

Neurogliaform interneurons exhibit different types of LTP (30). Indeed, theta burst stimulation (TBS) of the temporoammonic pathway produces robust LTP even with no induced somatic depolarization. This suggests that the temporoammonic synaptic inputs may be integrated by the postsynaptic dendrites to produce synaptic plasticity. To understand the rules governing TBS LTP induction in neurogliaform interneurons better, we analysed the LTP induction traces from Mercier et al. (30). We hypothesised that variability in LTP magnitude following theta-burst stimulation (TBS) in neurogliaform interneurons is better explained by differences in dendritic integration and action potential generation during the induction protocol than by baseline synaptic strength alone.

The TBS LTP induction protocol consisted of 3 sets of 10 theta bursts (**Figure 3a**). The membrane voltage was allowed to float in current clamp mode. The recorded cells exhibited a range of EPSP waveforms (**Figure 3b**). Some EPSPs decayed to baseline within the 200 ms period between stimulation bursts, some had a pronounced plateau-like waveform, and others were accompanied by action potentials. Likewise, the LTP magnitude varied across cells but seemed to be the most pronounced in cells firing action potentials as a result of the synaptic stimulation (**Figure 3c**).

**Figure 3.**
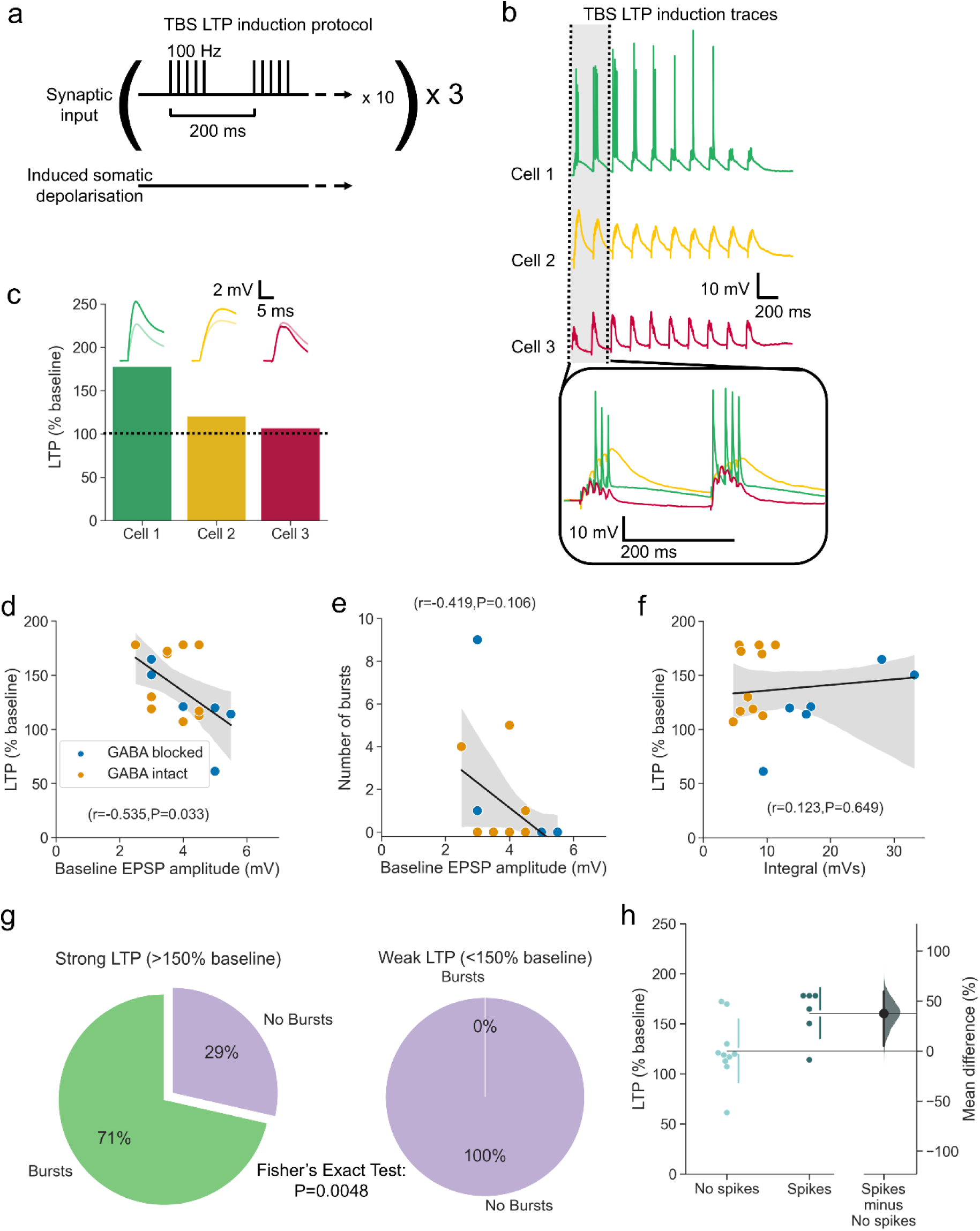
Dendritic integration may drive theta burst stimulation-induced LTP. **(a)** Overview of the theta burst stimulation (TBS) long-term potentiation (LTP) induction protocol. Synaptic input was elicited by electrically stimulating the temporoammonic pathway afferents. There was no induced somatic depolarization. **(b)** Example induction traces of the first set of bursts, with a zoomed in inset showcasing all three example traces. **(c)** Overview of the LTP (% baseline) exhibited by the example cells from (b). EPSPs before (light) and after (dark) LTP are shown correspondingly. **(d)** LTP plotted against baseline EPSP amplitude. In some recordings GABAergic transmission was blocked with picrotoxin (100 μM) and CGP55845 (1 μM) The color coding is conserved across panels (d) to (f). **(e)** Number of bursts plotted against baseline EPSP amplitude. **(f)** LTP plotted against integral. **(g)** Relationship between action potential bursts during LTP induction and the subsequent LTP. **(h)** LTP in cells with and without spikes during the induction. The mean difference between no spikes and spikes is 38% [95%CI 5%, 60%]. P=0.0242 (two-sided permutation t-test). The mean difference between the groups is shown in Gardner-Altman estimation plots as bootstrap sampling distributions. The mean difference is depicted as a black dot. The 95% confidence intervals are indicated by the ends of the vertical error bars. Shaded areas in panels (d-f) represent the 95% confidence interval. n = 16 cells from 6 animals.

We first asked whether the observed LTP magnitude was simply a product of a stronger baseline synaptic input, as might be expected from the principle of synaptic cooperativity (55). However, the LTP magnitude correlated negatively with the baseline EPSP (**Figure 3d**, Pearson: r=-0.535, P=0.033). There was also no evidence that a greater baseline EPSP amplitude led to more bursts of action potentials during TBS (**Figure 3e**, Spearman: r=0.419, P=0.106). There was also no correlation between the EPSP integral and LTP (**Figure 3f**, Pearson: r=0.123, P=0.649). In contrast, the majority of cells exhibiting strong LTP also fired bursts of action potentials, whilst no cells with weak LTP fired action potentials during TBS (**Figure 3g**, Fisher’s exact test: P=0.0048). When the cells were dichotomized based on whether there were any spikes during TBS, another clear effect emerged. LTP magnitude was 38% (P=0.0242, two-sided permutation t-test) greater in the cells that produced spikes.

These effects and associations suggest that a critical factor determining whether synaptic potentiation occurs is whether they fire any action potentials, and in particular whether they fire bursts of action potentials. Although bursts of action potentials are predictive of LTP, it cannot be said definitively whether they are a cause or consequence of the dendritic integration of synaptic input that causes LTP.

### Clustered TBS-like synaptic input results in greater calcium entry than dispersed synaptic input

In the *ex vivo* experiment from Mercier et al. (30) synaptic input was provided by electrically stimulating the temporoammonic pathway, meaning that every cell received a variable and indeterminate number of synaptic inputs. Different cells likely received synaptic inputs that exhibited different degrees of spatial clustering. A possible factor determining whether LTP is elicited is the local calcium concentration arising as a result of the synaptic input. We hypothesised that dendritic clustering of synaptic inputs during TBS leads to greater local calcium influx than spatially dispersed input, thereby providing a mechanistic basis for variability in LTP induction.

To test this hypothesis, a model cell was populated with 1 to 12 synapses which were then activated cumulatively as part of a 100 Hz TBS-like burst (**Figure 4a**). The dendritic NMDAR current at the sites of stimulation was captured and multiplied by the permeability of the NMDAR to calcium, estimated to be ∼7% (56–58). (The different driving force for calcium than for monovalent cations was ignored for simplicity.) The synapses were either clustered on dendrite 35 or distributed across 12 different dendrites as before (**Figure 4b-c**). The resulting estimated calcium current was substantially greater in the clustered condition compared to that in the dispersed condition (**Figure 4c-e**). Note that in these simulations we chose to focus on calcium flux and to ignore intracellular calcium dynamics, in order to avoid untested and unconstrained assumptions on calcium buffering in this cell type (59–63). This also puts the emphasis on local calcium sensors at the site of influx, in keeping with the mechanisms of induction of NMDAR-dependent LTP in principal neurons. We also assumed that individual synapses release on successive stimuli, thus ignoring probabilistic exocytosis and refractoriness. The conclusions of these simulations should therefore be taken as a qualitative guide to the conditions that maximize calcium influx in response to subthreshold stimulation.

**Figure 4.**
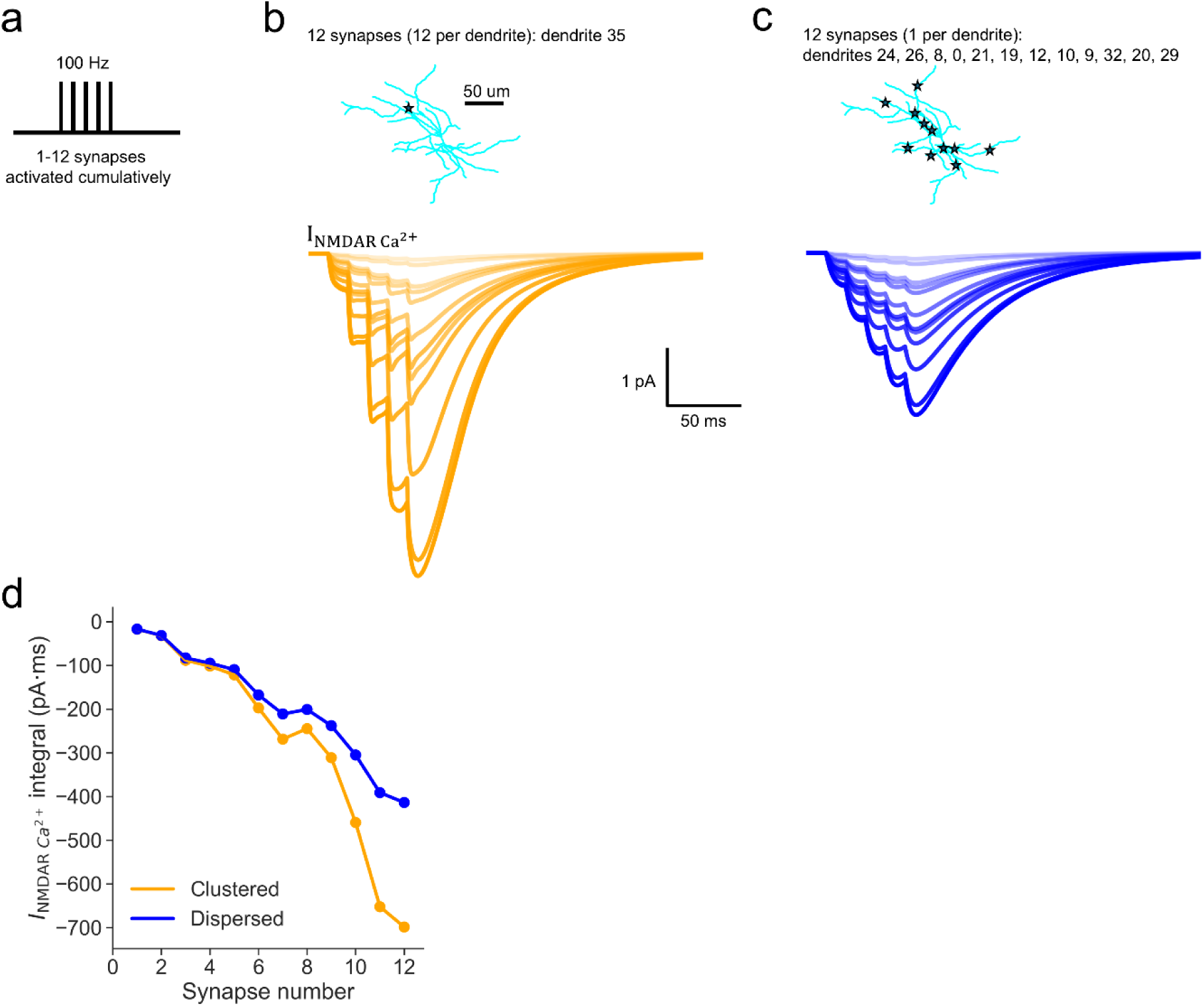
Clustered TBS-like synaptic input results in greater calcium entry via NMDARs than dispersed synaptic input. **(a)** Stimulation overview used in the simulation. Cumulatively increasing numbers of synapses were activated, 1 to 12. The synapses were activated simultaneously at 100 Hz to approximate the type of synaptic input delivered in the *ex vivo* TBS LTP experiment. **(b)** The calcium current conducted via the NMDAR was calculated by multiplying the NMDAR current (I) recorded during the simulation by a scaling factor (P) of 0.07, corresponding to the approximate NMDAR permeability to calcium. **(c)** Overview of synaptic input locations clustered on dendrite 35 and the corresponding NMDAR calcium current at the stimulated synapses. **(d)** As in (c) but with synaptic input locations dispersed across the dendritic tree. **(e)** NMDAR calcium current integral with increasing numbers of activated synapses in the clustered and dispersed conditions.

### Somatic action potentials can invade all dendrites whilst calcium currents attenuate rapidly with distance from soma

Action potentials have been shown to backpropagate into dendrites of principal cells (3, 4) and interneurons (64–66) including neurogliaform interneurons (67). We used the *in silico* model to characterize the spatiotemporal profile of action potential backpropagation in neurogliaform interneurons.

A single somatic action potential was elicited with a 1 ms somatic current injection in absence of synaptic input. This spike quickly propagated throughout the dendritic tree (Figure 5 a-c) and resulted in a very small VGCC-mediated calcium current that attenuated with distance from soma (**Figure 5j**). When a burst of 5 somatic action potentials was elicited at 100 Hz using 1 ms current injection steps instead, the depolarization propagated similarly throughout the dendritic tree (**Figure 5d-e**). However, the burst of action potentials evoked a far greater VGCC calcium current than the single spike (**Figure 5j**).

**Figure 5.**
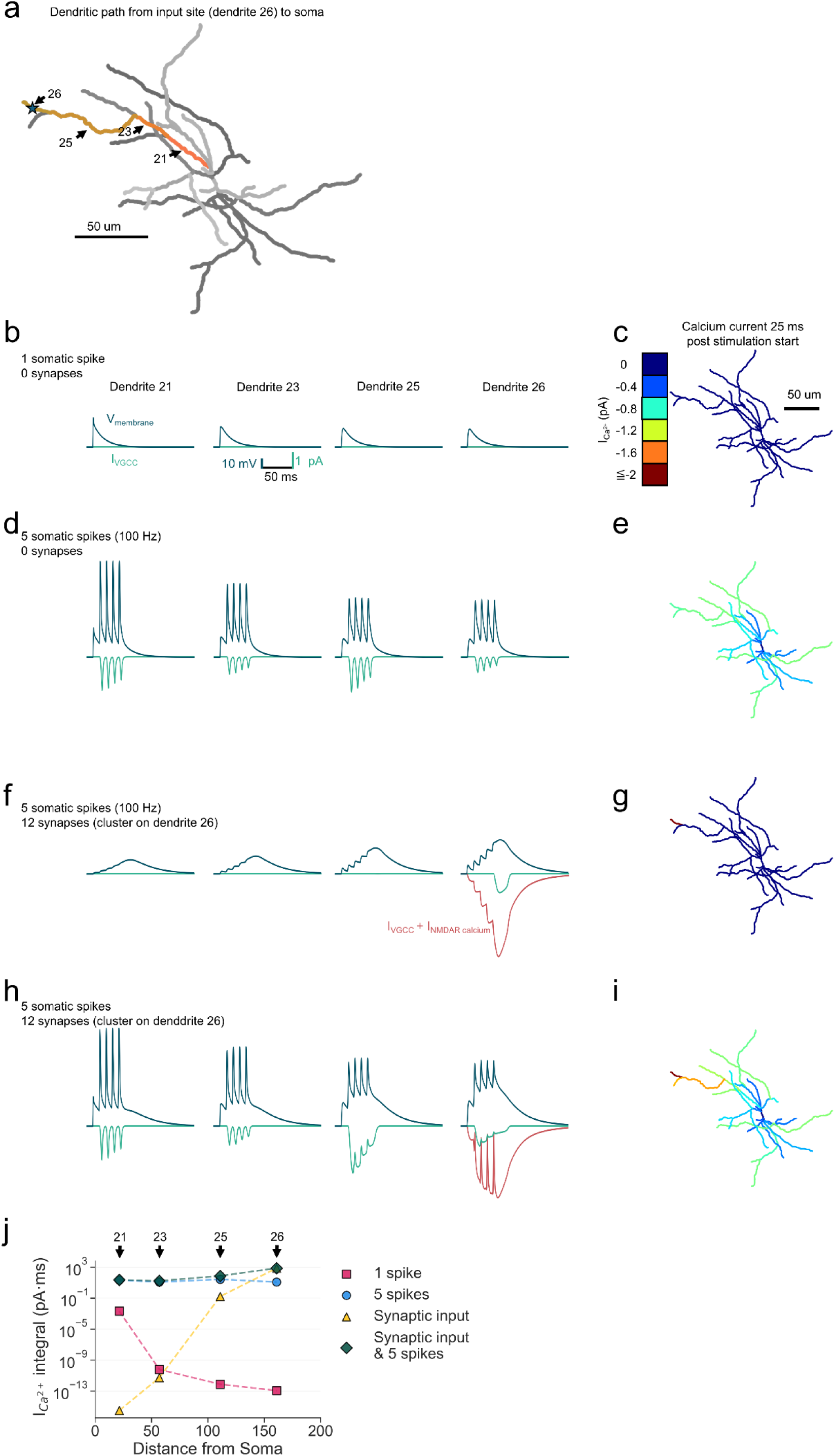
Somatic action potentials can invade distal dendrites and boost calcium currents arising from synaptic input. **(a)** A map of 4 dendrites (21, 23, 25, 26) connecting end-to-end, starting from the soma. Synaptic input locations on dendrite 26 indicated by stars. **(b)** Membrane voltage and VGCC calcium current following a single somatic action potential, measured at 4 different dendrites highlighted in (a). **(c)** The calcium current recorded in the simulated neuron, overlayed onto its morphology, 25 ms after stimulation start. **(d)** As in (b) but following a burst of 5 somatic action potentials delivered at 100 Hz. **(e)** As in (c) but corresponding to (d). **(f)** As in (b) but following a burst of synaptic input clustered on dendrite 26 and delivered at 100 Hz. NMDAR calcium current is also displayed. **(g)** As in (c) but corresponding to (f). **(h)** As in (f) but following a burst of 5 somatic action potentials delivered at 100 Hz as well as a concurrent synaptic input clustered on dendrite 26 and delivered at 100 Hz. **(i)** As in (c) but corresponding to (h). **(j)** Integral of total calcium current conducted by VGCCs and NMDARs, with increasing distance from the soma and expressed as an absolute value. Numbers above the plot indicate the dendrite measured.

Notably, the VGCC calcium current produced in both cases, arising from a single action potential and from a burst of action potentials, attenuated with distance from the soma. This can be seen readily when focusing on a series of parent-child dendrites, dendrites 21, 23, 25, 26 (**Figure 5a, j**). Action potential amplitudes attenuated as the signal propagated down the dendrites but even the most distal compartment, dendrite 26, was depolarized eventually (**Figure 5b-e**). The elicited VGCC calcium current, on the other hand, progressively collapsed toward 0 as the distance from the soma increased (**Figure 5b-e, j**). This suggests that the backpropagating voltage and calcium signals dissociate across space when driven by somatic action potentials alone.

### Concurrent synaptic input and somatic action potentials boost local membrane depolarization and calcium influx to overcome the distance-dependent attenuation of calcium current

How does synaptic input interact with action potentials in the context of calcium currents? To investigate this question, we repeated the simulations without action potentials and added twelve synapses clustered onto the distal dendrite 26.

Under these conditions, the local voltage dynamics appeared to be qualitatively similar in the distal dendrites and at or close to the site of synaptic input (**Figure 5f**). However, total calcium current dynamics (the sum of NMDAR and VGCC calcium currents) told a different story. Whilst there was a substantial VGCC- and NMDAR-mediated calcium current in the stimulated dendrite 26, the depolarization in dendrites 25, 23, and 21 was insufficient to produce any appreciable influx of calcium (**Figure 5g**). Indeed, with increasing distance from the stimulated dendrite, the calcium current was profoundly attenuated (**Figure 5j**).

We next simulated coincident synaptic input and somatic action potentials to ask whether this overcomes the spatial attenuation observed during backpropagating spikes or synaptic input alone. Simultaneous somatic action potentials and synaptic input produced large depolarizations across all compartments (**Figure 5h**). A large total calcium current was also elicited in many compartments (**Figure 5h-i**). Following the same parent-child series of dendrites 21 to 26, it is clear that both the voltage and calcium signals remained relatively large throughout (**Figure 5h-j**). When both synaptic input and induced somatic spikes coincided in time, neither the voltage nor the calcium signals attenuated with distance in relative terms (**Figure 5h-j**). This result suggests that somatic action potentials interact with and facilitate the forward-propagation of the synaptic calcium signal to the soma.

### Spike backpropagation attenuates with distance from soma and dendrite order ex vivo

To characterize calcium influx elicited by backpropagating somatic action potentials in neurogliaform interneurons and to test our *in silico* predictions on calcium entry due to backpropagating spikes we conducted a set of *ex vivo* experiments. NDNF-positive (NDNF+) neurogliaform interneurons were studied in the hippocampal CA1 stratum lacunosum moleculare in acute slices from *Ndnf^cre^*or *Ndnf^cre/cre^* mice injected with AAV2/9-mDLX-FLEX-mCherry virus. NDNF+ neurogliaform interneurons were patch-clamped, filled with Alexa-594 and the fluorescent calcium-sensor Fluo-4, and stimulated by inducing 0 to 5 somatic spikes (**Figure 6a-b**). Fluo-4 calcium fluorescence was measured via linescans on sections of dendrites at varying distances from the soma (**Figure 6c**).

**Figure 6.**
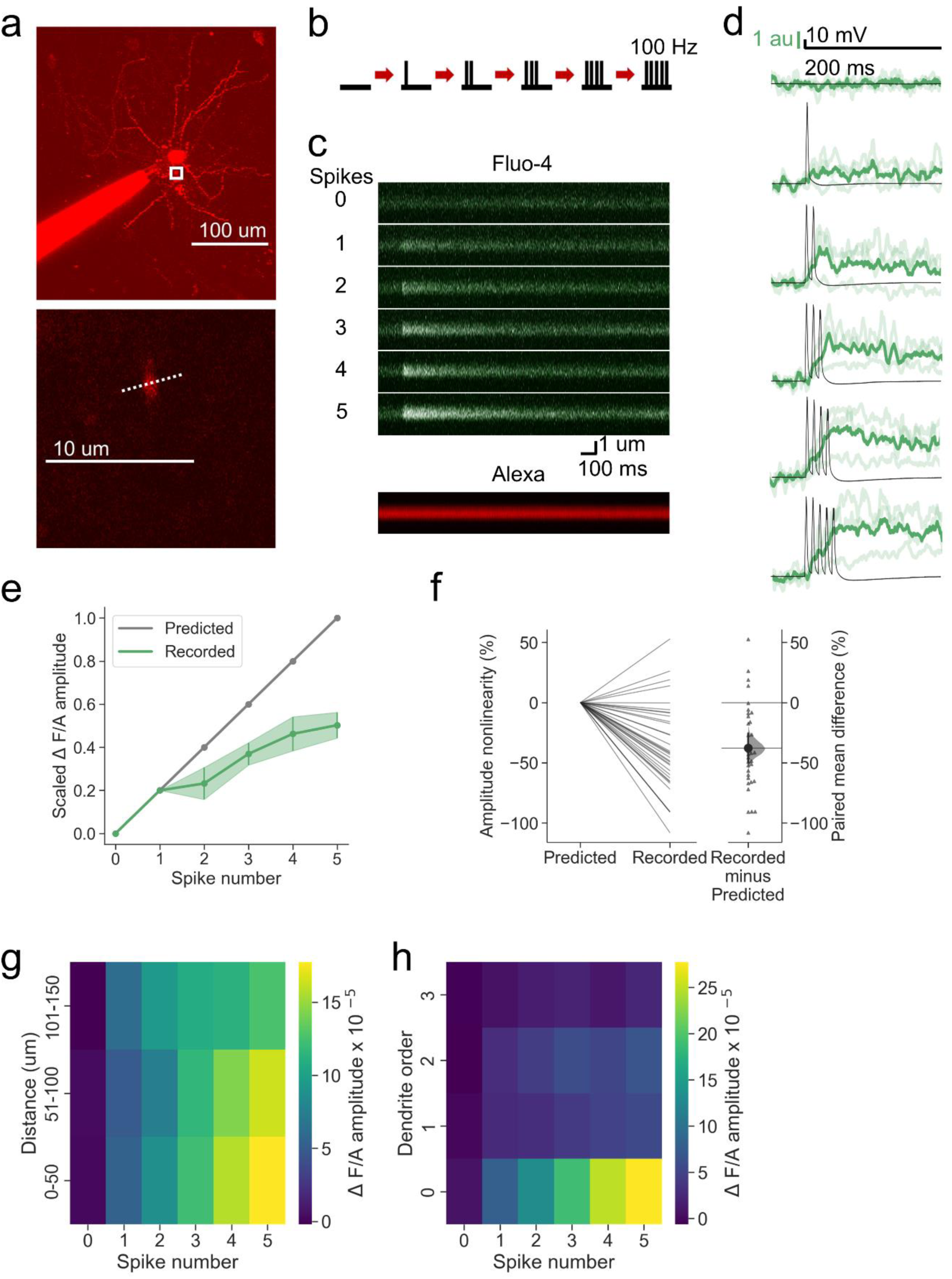
Spike backpropagation attenuates with distance from soma and dendrite order *ex vivo*. **(a)** A representative neurogliaform interneuron patch-filled with Alexa-594. Inset indicates dendrite where calcium fluorescence was recorded. Dashed line marks the position of the linescan. **(b)** Stimulation protocol. Increasing numbers of spikes were generated, from 0 to 5 (100 Hz). **(c)** Linescans showing calcium Fluo-4 and Alexa-594 fluorescence. **(d)** Overlaid somatic voltage and dendritic calcium fluorescence traces. Dendritic calcium traces are shown as a ratio of Fluo-4 to Alexa-594 (arbitrary units). A mean of 3 trials is shown as darker color traces and the individual trials are shown as lighter color traces. **(e)** Scaled change in calcium fluorescence amplitude as a function of action potential spike number. SEM indicated by vertical bars. **(f)** The paired mean difference between predicted and recorded is -38% [95%CI -49%, - 25%]. P<0.001 (two-sided permutation t-test compared to 0%). For the slopegraph and Gardner-Altman estimation plot, paired observations are connected by lines. The paired mean difference between the predicted and recorded groups is shown in Gardner-Altman estimation plot as a bootstrap sampling distribution. The mean difference is depicted as a black dot. The 95% confidence intervals are indicated by the ends of the vertical error bars. **(g)** Heatmap of calcium-fluorescence as a function of distance from soma and spike number. **(h)** Heatmap of calcium-fluorescence as a function of dendrite order and spike number. n = 34 dendrites in 19 cells from 14 animals.

The amplitude of the calcium fluorescence transients in the dendrites exhibited a positive dependence on the number of somatic spikes (**Figure 6c-e**). This increase in calcium fluorescence was however sublinear (**Figure 6f**), potentially due to the absence of concurrent synaptic input to activate regenerative NMDAR conductances.

Dendritic calcium fluorescence was observed at relatively short (0-50 µm), intermediate (51-100 µm), and long (101-150 µm) distances from the soma (**Figure 6g**). There was a small and non-significant decrease in calcium fluorescence as the distance from soma increased (Pearson: r<0.01, P=0.96). A more striking contrast emerged when the same data were arranged by dendrite order (**Figure 6h**). Although calcium-dependent fluorescence transients were seen in dendrites of all orders tested (0–3), the fluorescence amplitude was the greatest in dendrites of the zeroth order (Spearman: r=-0.37, P=0.03).

Taken together, these data align with the modelling predictions, namely that somatic action potentials invade even relatively distant compartments and that backpropagating calcium signals attenuate with increasing distance from the soma.

### Coincident somatic spikes increase uEPSP amplitude and attenuate supralinearity

In addition to TBS, a spike-pairing protocol (synaptic input followed by an induced somatic action potential) has also been shown to induce synaptic potentiation in neurogliaform interneurons (30). Hippocampal principal neurons exhibit a wide variety of spike timing windows in the context of synaptic plasticity (68). We took advantage of the *in silico* model to investigate the optimal temporal alignment of synaptic input and somatic action potential. We hypothesised that the relative timing between synaptic input and somatic action potentials determines the degree of dendritic supralinearity and calcium influx, with different temporal orders producing distinct integration regimes.

The same 12 synapses were activated on dendrite 35, and in addition a somatic action potential was induced coincidently, 25 ms before or after the onset of synaptic stimulation. When synaptic input coincided with the somatic spike, a rapid depolarization spread throughout the dendritic tree (**Figure 7a**). The maximal uEPSP amplitude was greater than that generated from synaptic input alone (**Figure 7b**). The median degree of supralinearity was however lower than seen with equivalent synaptic stimulation without a somatic spike (119% vs 125%) (**Figure 7g**). The maximum degree of supralinearity was seen in the stimulated dendrite 35 and was also lower than after equivalent synaptic stimulation with no somatic spike (134% vs 144%).

**Figure 7.**
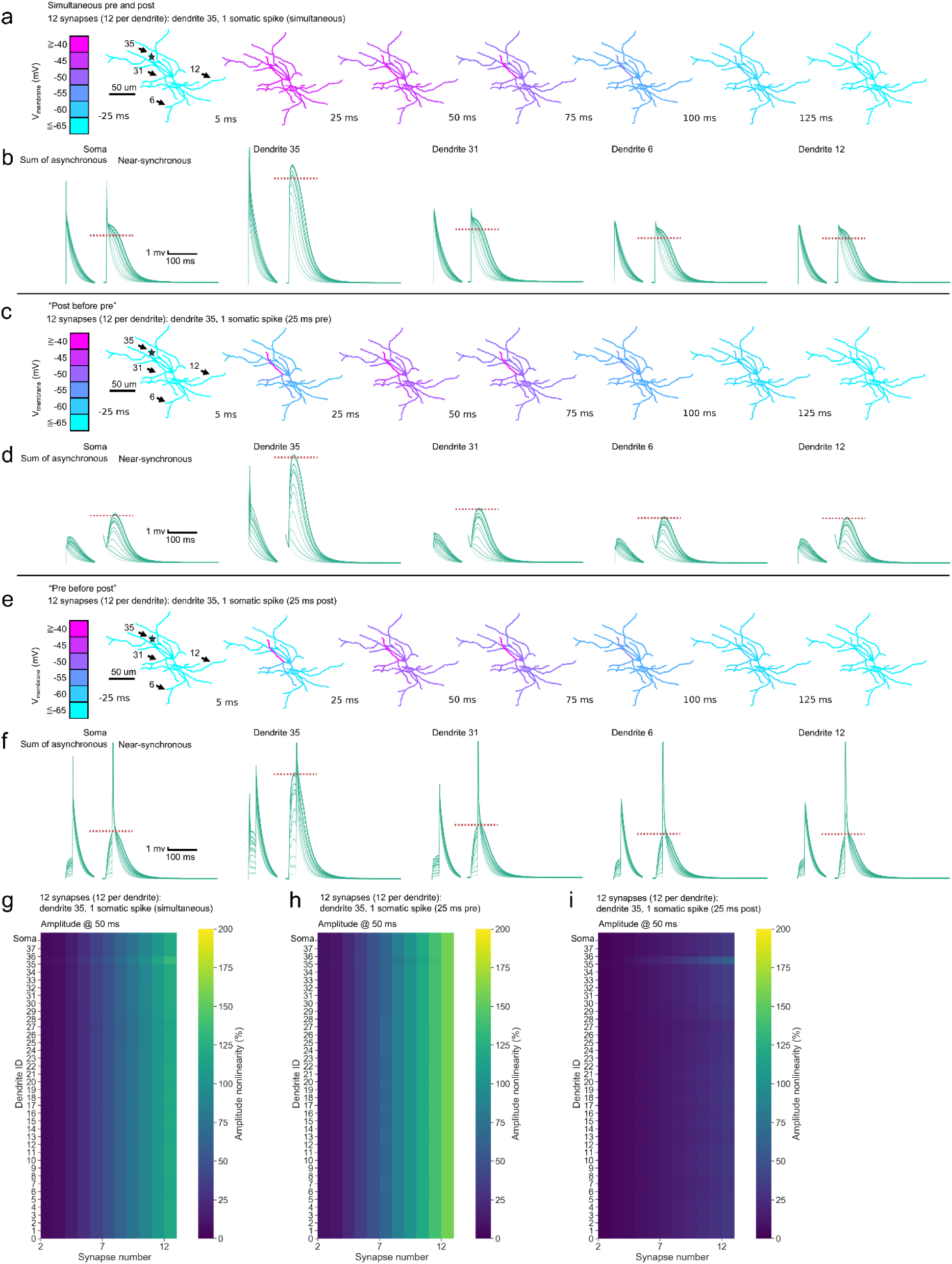
Coincident somatic spikes increase uEPSP amplitude and attenuate supralinearity. **(a)** Simulated clustered synaptic input with 1 somatic spike (coincident). 12 synapses were activated on dendrite 35. Voltage snapshots at 25 ms before stimulation and 5-125 ms after stimulation onset overlaid onto the neuronal architecture. Stimulation site indicated by the star. Dendrites 35, 31, 6, 12 indicated with arrows have their uEPSPs displayed as traces below. **(b)** Sum of the asynchronous s and near-synchronous uEPSPs at the soma and dendrites 35, 31, 6, 12. The maximum amplitude of the near-synchronous response without a spike (from Figure 2) is indicated by a red dashed line. **(c)** Simulated clustered synaptic input with 1 somatic spike (25 ms pre onset of synaptic stimulation). Data displayed as in (b). **(d)** Sum of the asynchronous s and near-synchronous uEPSPs at the soma and dendrites 35, 31, 6, 12. The action potential that preceded the first uEPSP used in the sum is only partially displayed (decay visible, not spike). **(e)** Simulated clustered synaptic input with 1 somatic spike (25 ms post onset of synaptic stimulation). Data displayed as in (b). **(f)** Sum of the asynchronous s and near-synchronous uEPSPs at the soma and dendrites 35, 31, 6, 12. **(g)** Heatmap of amplitude nonlinearity for every dendrite and the soma. Corresponds to (a, b). **(h)** Heatmap of amplitude nonlinearity for every dendrite and the soma. Corresponds to (c, d). **(i)** Heatmap of amplitude nonlinearity for every dendrite and the soma. Corresponds to (e, f). Amplitude was measured at 50 ms post stimulus onset in (g-i).

When the synaptic input was preceded by a somatic spike (‘post before pre’), the spike backpropagated through the dendritic tree, depolarizing all compartments by ∼10mV before the onset of the synaptic stimulation. The synaptic stimulation then elicited an uEPSP that in turn invaded other compartments (**Figure 7c**). The maximal uEPSP amplitude was greater than that evoked by synaptic input alone (**Figure 7d**), similar to when the synaptic input somatic spike were coincident. The median degree of supralinearity was greater than seen with equivalent stimulation without a somatic spike (157% vs 125%) (**Figure 7h**). The maximum degree of supralinearity was exhibited by the stimulated dendrite 35 (157%). Notably both the median and maximum degrees of supralinearity under this paradigm were greater than when synaptic input and a somatic spike were completely coincident. The magnitude of supralinearity across dendrites was also notably similar across dendrites.

When the somatic spike was elicited 25 ms after the onset of synaptic input (‘pre before post’), the dendritic integration phenotype changed dramatically. The maximum uEPSP amplitude was approximately equal to that evoked by the synaptic input alone (**Figure 7e-f**). The median degree of supralinearity was profoundly reduced compared to that seen with equivalent stimulation without a somatic spike (32% vs 125%) (**Figure 7h**). The median degree of supralinearity was also substantially lower than with a coincident somatic spike (32% vs 119%). The maximum degree of supralinearity was exhibited by the stimulated dendrite 35 (60%). Both the median and maximum degrees of supralinearity achieved under this configuration were even lower than when 12 dispersed synapses were activated (median: 55%, maximum: 60%) (**Figure 2c-d**).

### Calcium current at the site of synaptic input is sensitive to somatic action potential timing during simulated spike pairing induction

The simulations in Figure 7 reveal an unexpected pattern whereby both the absolute dendritic depolarization and the degree of supralinearity are reduced under the ‘pre before post’ condition when compared to the other two conditions. How does this relate to calcium influx? It could be hypothesized that in addition to there being different magnitudes of voltage supralinearity across dendrites in response to the relative timings of the synaptic input and action potentials, there could be altered calcium dynamics as well.

We therefore quantified calcium flux in the spike pairing simulations. All three conditions had prominent NMDAR calcium currents (**Figure 8a-c**). VGCCs were engaged in all three conditions as well. Total calcium current was similar whether the action potential was coincident with the synaptic input or whether the action potential preceded synaptic input, “post before pre” (**Figure 8d-e, g**). On the other hand, and in agreement with the voltage nonlinearity results, when the action potential followed synaptic input, “pre before post”, calcium currents were attenuated (**Figure 8f-g**). The total calcium current integral was the greatest under the simultaneous condition, when the action potential coincided with synaptic input (**Figure 8g**). The relative timings of the synaptic input and the action potential changed the proportion of the total calcium current that was either NMDAR or VGCC mediated.

**Figure 8.**
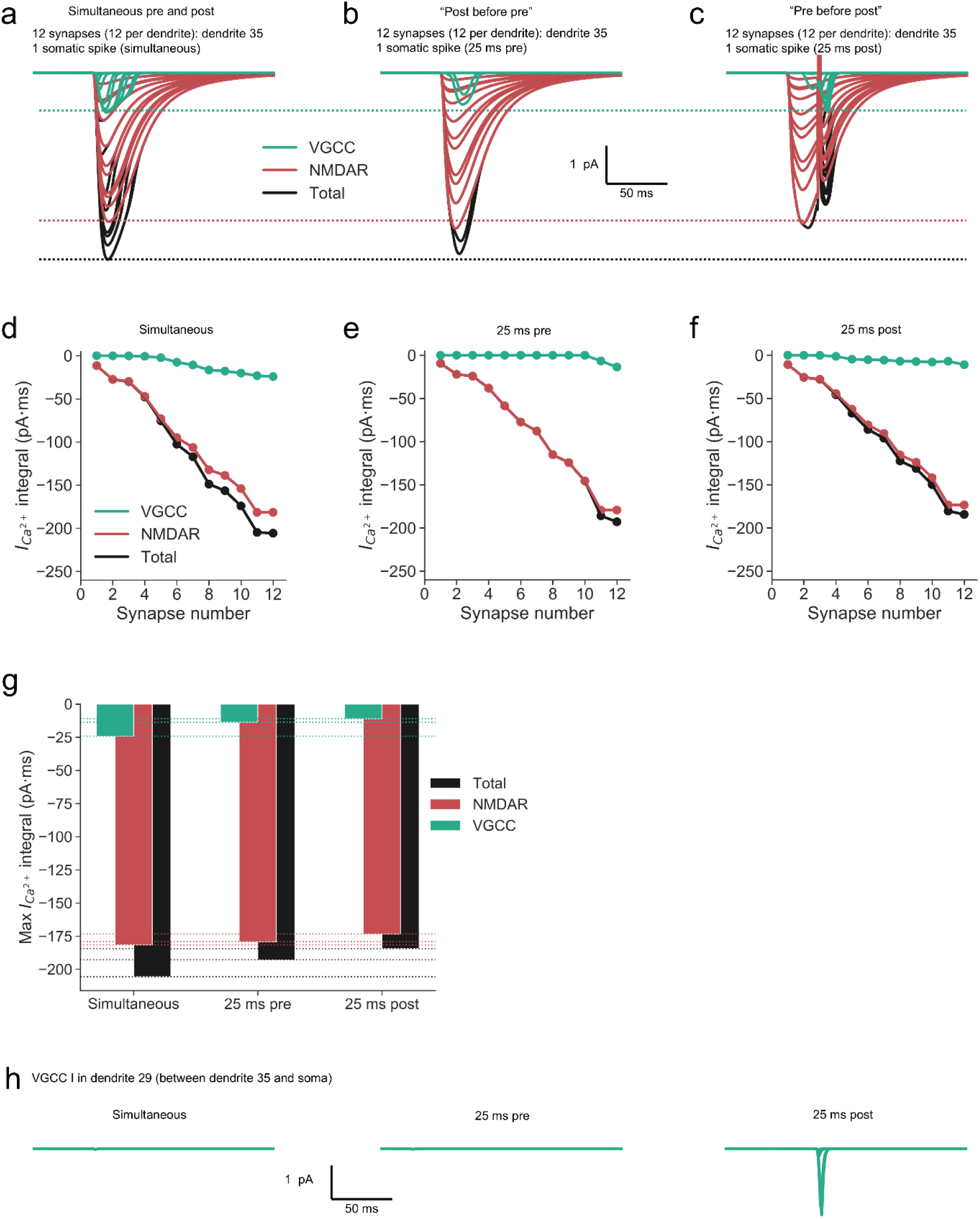
Calcium current at the site of synaptic input is insensitive to somatic action potential timing during simulated spike-pairing induction. Calcium current traces from dendrite 35 following cumulative synaptic input and a somatic spike elicited simultaneously with synaptic input **(a)**, 25 ms before synaptic input **(b)**, and 25 ms after synaptic input **(c)**. Dashed lines drawn to facilitate visual comparison. Calcium current integral with increasing synapse numbers activated **(d-f)**, corresponding to the above traces (a-c). **(g)** Summary bar plot comparing maximum calcium current integral across conditions. **(h)** VGCC current in dendrite 29, which is between dendrite 35 that received synaptic input and the soma.

Overall, under the parameters tested, the calcium influx at the stimulated dendrite was sensitive to the differences in input timing. The input timing also impacted the VGCC current recorded in dendrite 29, which is between dendrite 35 (which received synaptic input) and the soma (**Figure 8h**). In dendrite 29 the greatest calcium flux was recorded when the synaptic inputs preceded the action potential. The relative timing of synaptic input and action potential firing is thus critically important for VGCC activation at sites where orthodromic synaptic and antidromic somatic depolarizations collide.

## Discussion

The main conclusion of this study is that, despite the absence of spines and involvement of sodium channels in supralinear summation, coincident synaptic activity and backpropagating action potentials elicit robust NMDAR-mediated calcium influx in neurogliaform interneurons, a necessary condition for the induction of an important form of LTP. There are however some subtle differences, notably in that depolarization and calcium signalling readily spread through the dendritic tree, and that they are sensitive to the precise temporal order of presynaptic activation and postsynaptic action potentials.

Synaptic inputs clustered on nearby dendritic sites have been reported *in vivo* (51, 52). On the other hand, some reports have indicated that synaptic inputs may be dispersed across the dendritic architecture (53, 54). Our results show that while both clustered and dispersed synaptic inputs elicit supralinear responses in neurogliaform interneurons, clustered inputs induce more pronounced local depolarizations. This presents the possibility that in this specific cell type supralinear dendritic integration may play a role regardless of the spatial input arrangement. Indeed, the small size of neurogliaform interneurons challenges the ability to compartmentalize postsynaptic depolarizations. They may have evolved to primarily detect the temporal coherence of their inputs – with the spatial arrangement as an additional tuning knob available to the cell and circuit. They are also inter-connected and signal relatively diffusely via ‘volume transmission’, extending the principle that time matters more than space.

Our *in silico* simulations led to the prediction that somatic action potentials can backpropagate into the dendrites of hippocampal neurogliaform interneurons. We confirmed this prediction *ex vivo* with calcium imaging. Even single spikes can backpropagate into dendrites up to ∼150 µm away, both *in silico* and *ex vivo*. We found that calcium fluorescence arising from backpropagating action potentials decreases with increasing distance from the soma and dendrite order. This phenotype is generally in line with expectation in terms of backpropagation (69). The possibility that all dendrites may become depolarized due to backpropagating spikes aligns with recent *in vivo* voltage imaging findings that showed little sign of electrical compartmentalization across dendrites (8).

Neurogliaform interneurons exhibit both NMDAR-dependent and NMDAR-independent LTP, suggesting the presence of multiple mechanisms to create a lasting trace of synaptic input (30). We found that in a theta burst protocol where the cells were allowed to spike and no action potentials were artificially induced, some cells produced bursts which were in turn associated with strong LTP. Further simulations revealed that clustered inputs could produce greater local calcium influx compared to dispersed inputs. We also found that coincident synaptic input and somatic spikes would result in calcium influx not only at the stimulated sites but also along the dendrites in between. Coincident synaptic input and backpropagating action potentials therefore may enhance EPSP amplitude and increase calcium flux along the dendrite, enabling voltage and calcium signal propagation and thereby contributing to synaptic plasticity when occurring coincidently. The spread of calcium signalling provides an explanation for a small but significant potentiation of a control pathway that was not active during a pairing protocol used to induce LTP (30).

In the context of the spike-timing LTP protocol, we also predicted greater calcium flux at the stimulated synapse when the action potential was coincident with the synaptic input or when it preceded synaptic input by 25 ms. Somewhat unexpectedly, voltage supralinearity was attenuated when the somatic action potential occurred 25 ms after the synaptic input. This ‘pre before post“ sequence is conventionally held to be optimal to induce LTP (5, 70), although the experimental evidence from principal cells is far from unanimous (68). A full exploration of the implications of these differences would require simulation or experimental manipulation of calcium buffering in the different compartments.

Our findings highlight the complex interplay among synaptic input, morphology, signal bidirectionality, and dendritic supralinearity in neurogliaform interneurons. The computational modelling provided key predictions, some of which were supported by experimental data, while others warrant further investigation. Neurogliaform interneurons appear to have the capacity to interpret temporally coherent converging synaptic inputs, whether clustered or dispersed, and to interpret the two-way street that is forward- and backpropagating voltage and calcium signals.

---

**Supplementary Figure S1.**
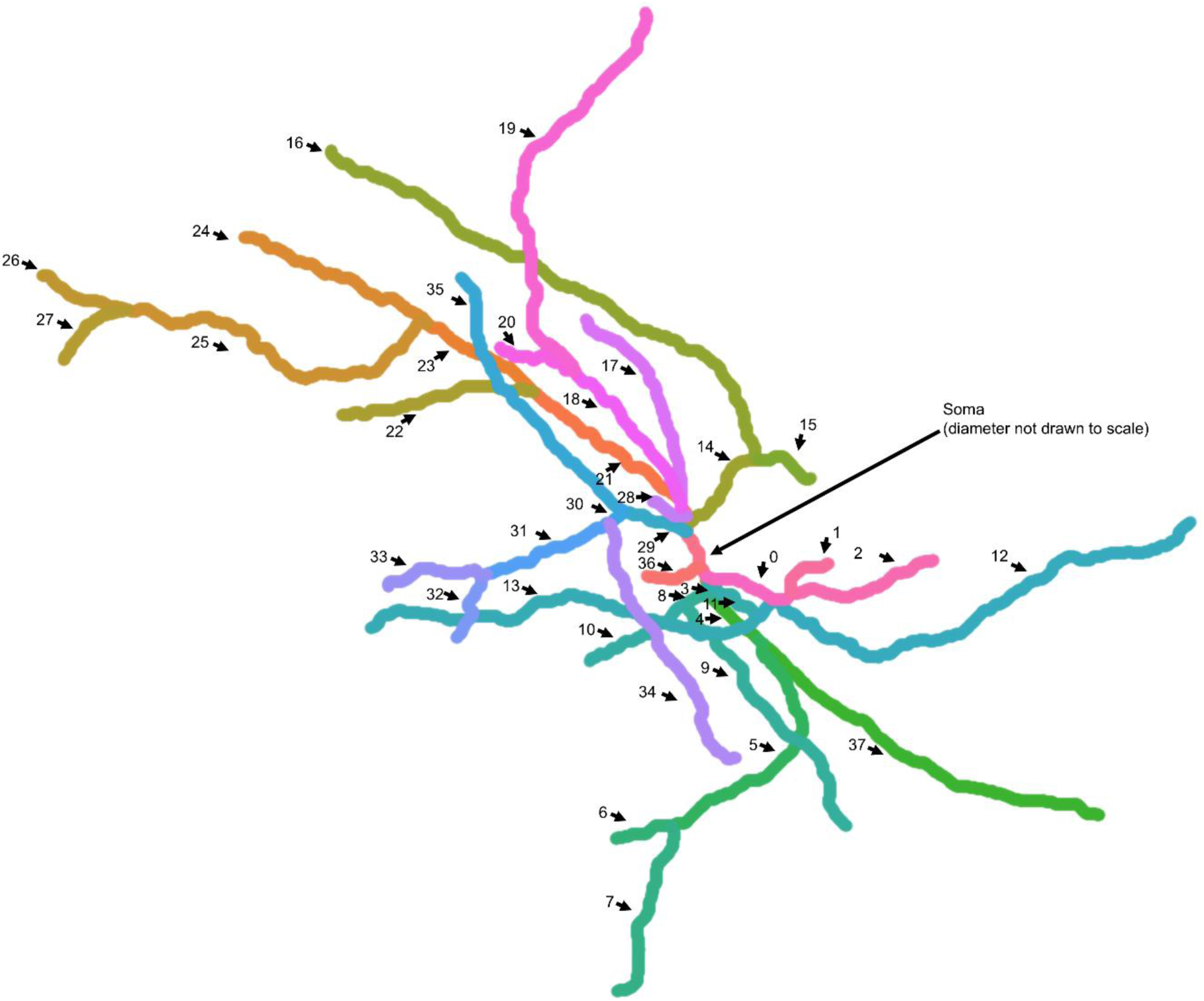
Map of the multi-compartment neurogliaform model. Dendrites 0-37 are identified by arrows. The position of the soma is also indicated.

**Supplementary Figure S2.**
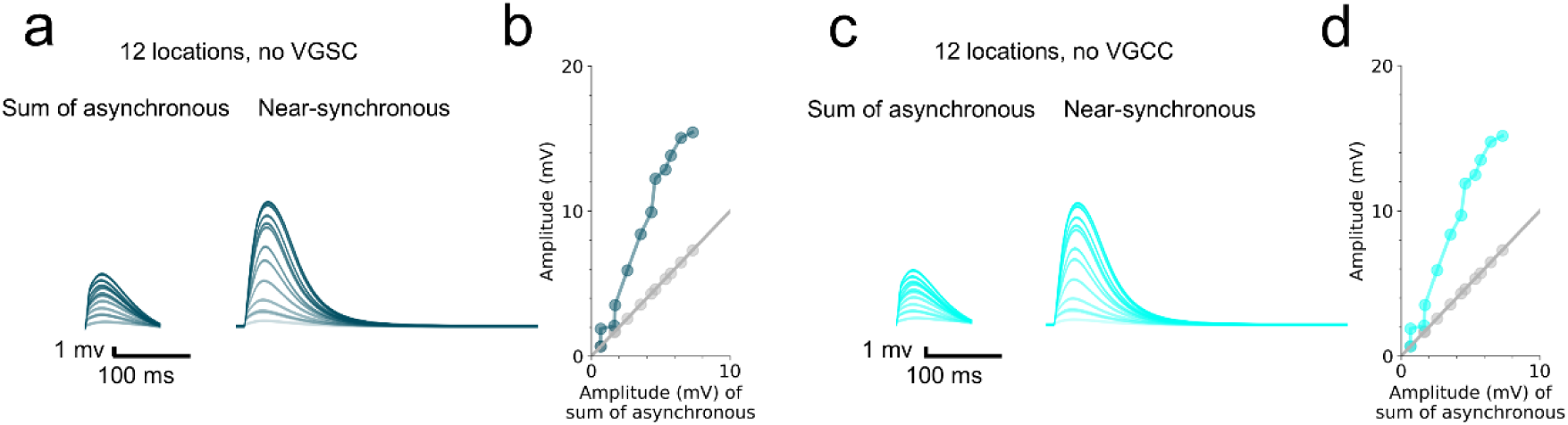
Effect of removing VGSCs or VGCCs from the multi-compartment model. **(a)** Simulated somatic voltage traces across asynchronous and near-synchronous conditions with VGSCs removed. **(b)** Amplitude of the simulated near-synchronous response plotted against the amplitude of the sum as the number of activated synapses was increased. The line of identity is represented by a grey line. **(c)** As in (a) but with L-type VGCC removed. **(d)** As in (b) but with L-type VGCC removed.

**Supplementary Figure S3.**
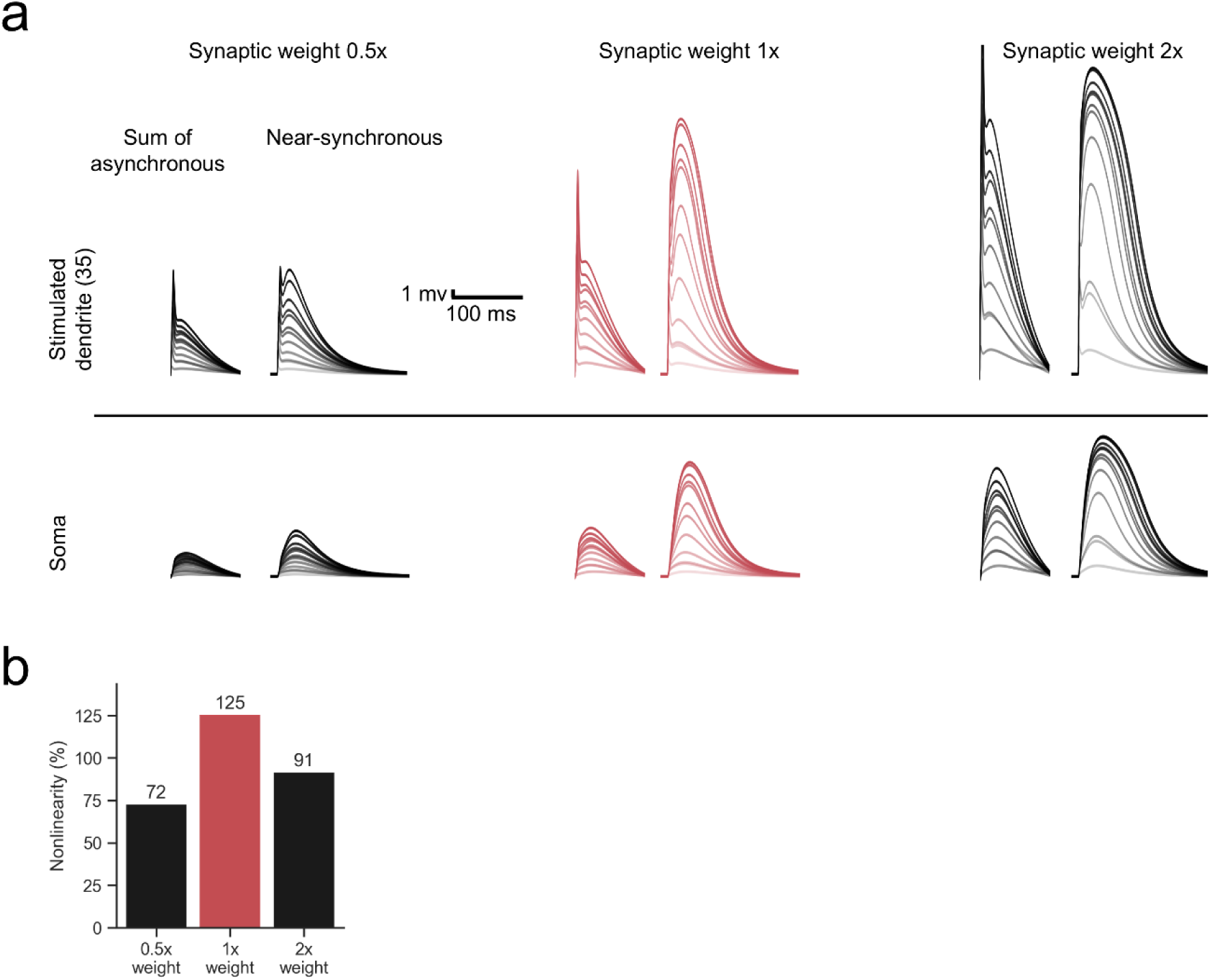
Changing the synaptic weights alters the magnitude of uEPSP nonlinearity at the stimulated dendrite and the soma. **(a)** Families of uEPSP traces following simulated clustered synaptic input on dendrite 35 and the soma. The sum of asynchronous uEPSPs and the near-synchronous uEPSPs are displayed for every condition. Synaptic weights of the glutamatergic synapses (both AMPAR and NMDAR components together) were scaled by a factor of 0.5 to 2 in different simulations. **(b)** Calculated nonlinearity across the different synaptic weights tested.

**Supplementary Figure S4.**
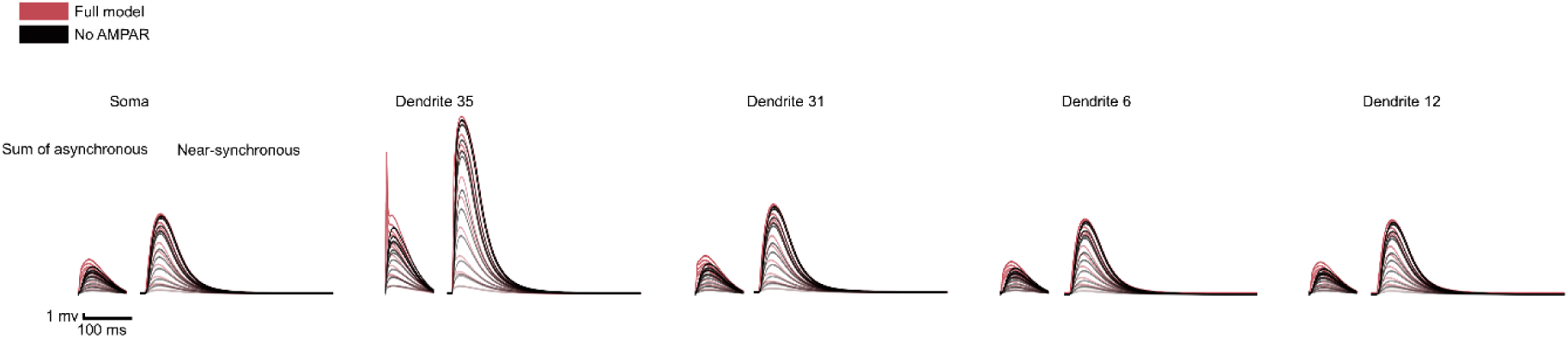
Effect of removing AMPAR from the multi-compartment model. Simulated clustered synaptic input delivered onto dendrite 35. 12 synapses were activated either with NMDARs modelled (full model) or with NMDARs removed. Traces of the sum of asynchronous and near-synchronous uEPSPs at the soma and dendrites 35, 31, 6, 12 are displayed.

**Supplementary Figure S5.**
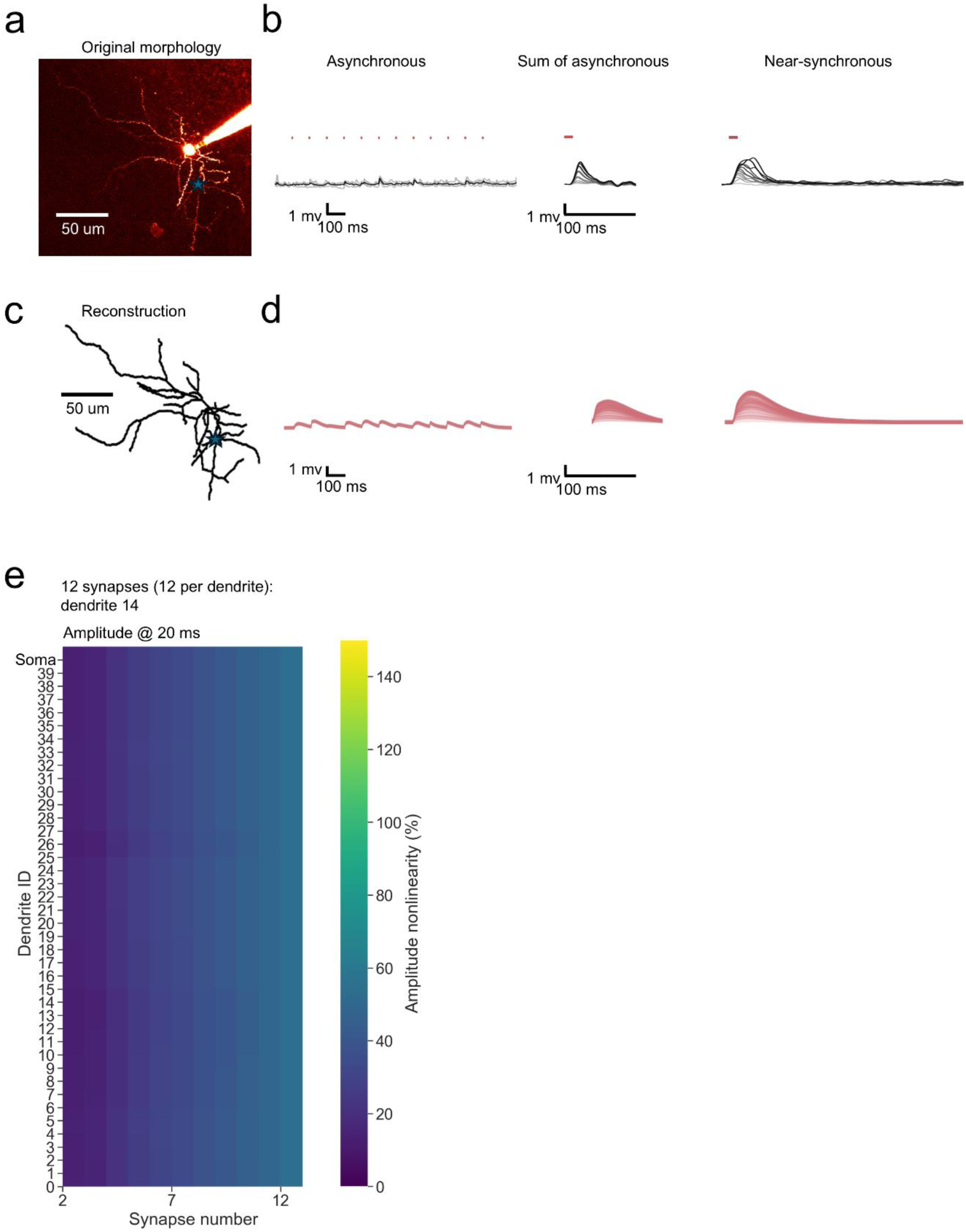
The *in silico* model reproduces the key *ex vivo* experimental finding of uEPSP amplitude supralinearity in response to near-synchronous synaptic input. **(a)** A second neurogliaform interneuron patch-filled with the inert dye Alexa-594. **(b)** Somatic voltage uEPSP traces across asynchronous and near-synchronous conditions. The sum of asynchronous uEPSPs is also displayed. Laser uncaging light stimuli are indicated by red dots above the traces. uEPSPs were elicited in the asynchronous condition with light pulses separated by an interval of 100.32ms (left; grey traces: individual sweeps; black traces average). The sum of asynchronous uEPSP traces were calculated by aligning and summing an increasing number of asynchronously evoked uEPSPs. Near-synchronous uEPSPs were elicited with light pulses separated by an interval of 0.32ms (right). The families of traces for the sum of asynchronous and near-synchronous uEPSPs are shown as light to dark grey traces, indicating an increase in the number of uncaging loci from 1 to 12. **(c)** The neuron shown in (a) was reconstructed and segmented into a multi-compartment model populated with passive and active properties. All parameters were set as in the main model presented in main figures 1-8. The synaptic weight was adjusted to approximate the sum of asynchronous traces amplitudes. **(d)** Simulated somatic voltage traces across asynchronous and near-synchronous conditions. Displayed as in (b). **(e)** Heatmap of amplitude nonlinearity for every dendrite and the soma. Corresponds to (c, d).

**Supplementary Figure S6.**
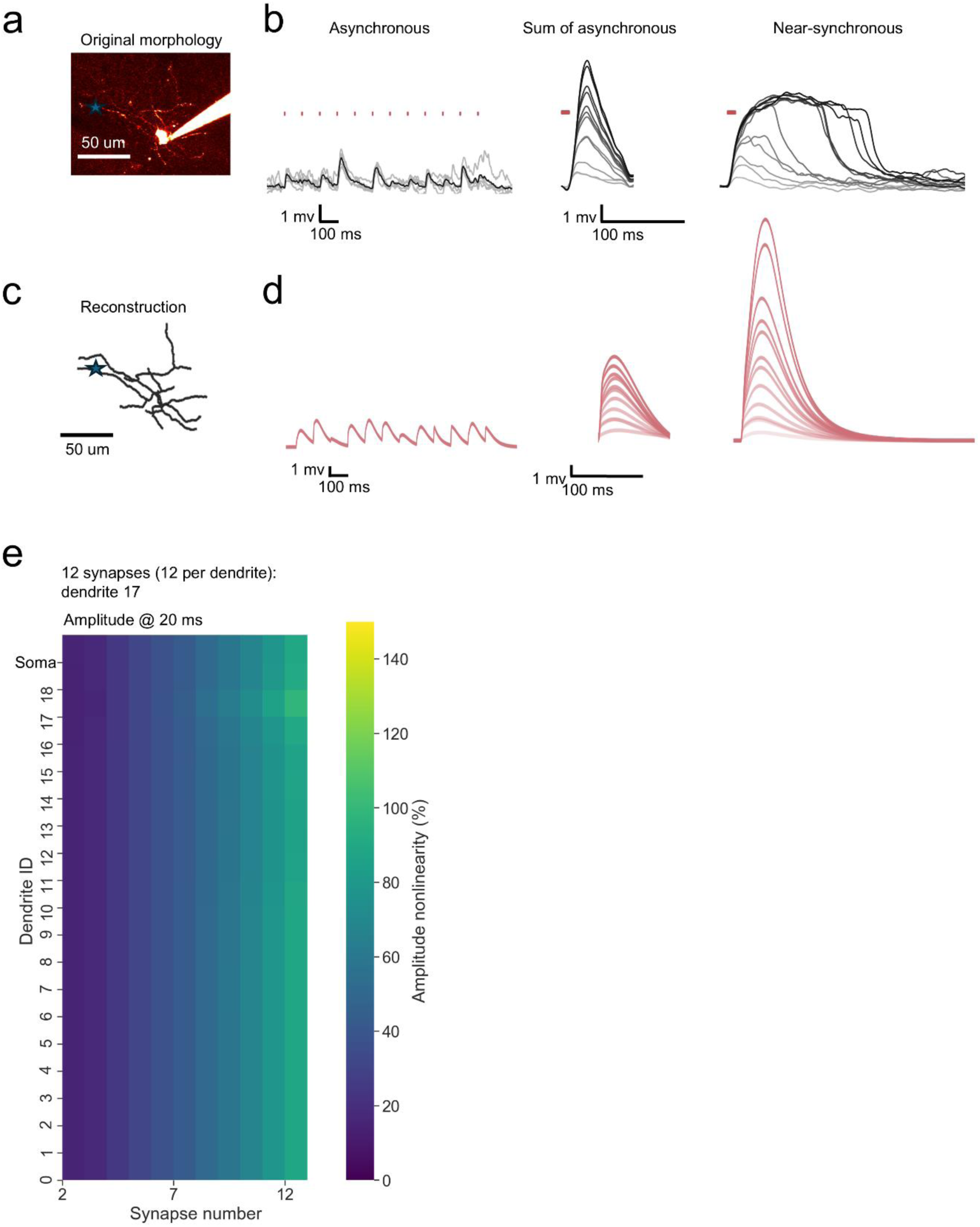
The *in silico* model reproduces the key *ex vivo* experimental finding of uEPSP amplitude supralinearity in response to near-synchronous synaptic input but does not produce long-lasting voltage “plateaus”. **(a)** A third neurogliaform interneuron patch-filled with the inert dye Alexa-594. **(b)** Somatic voltage uEPSP traces across asynchronous and near-synchronous conditions. The sum of asynchronous uEPSPs is also displayed. Laser uncaging light stimuli are indicated by red dots above the traces. uEPSPs were elicited in the asynchronous condition with light pulses separated by an interval of 100.32ms (left; grey traces: individual sweeps; black traces average). The sum of asynchronous uEPSP traces were calculated by aligning and summing an increasing number of asynchronously evoked uEPSPs. Near-synchronous uEPSPs were elicited with light pulses separated by an interval of 0.32ms (right). The families of traces for the sum of asynchronous and near-synchronous uEPSPs are shown as light to dark grey traces, indicating an increase in the number of uncaging loci from 1 to 12. **(c)** The neuron shown in (a) was reconstructed and segmented into a multi-compartment model populated with passive and active properties. All parameters were set as in the main model presented in main figures 1-8. The synaptic weight was adjusted to approximate the sum of asynchronous traces amplitudes. **(d)** Simulated somatic voltage traces across asynchronous and near-synchronous conditions. Displayed as in (b). **(e)** Heatmap of amplitude nonlinearity for every dendrite and the soma. Corresponds to (c, d).

**Supplementary Figure S7.**
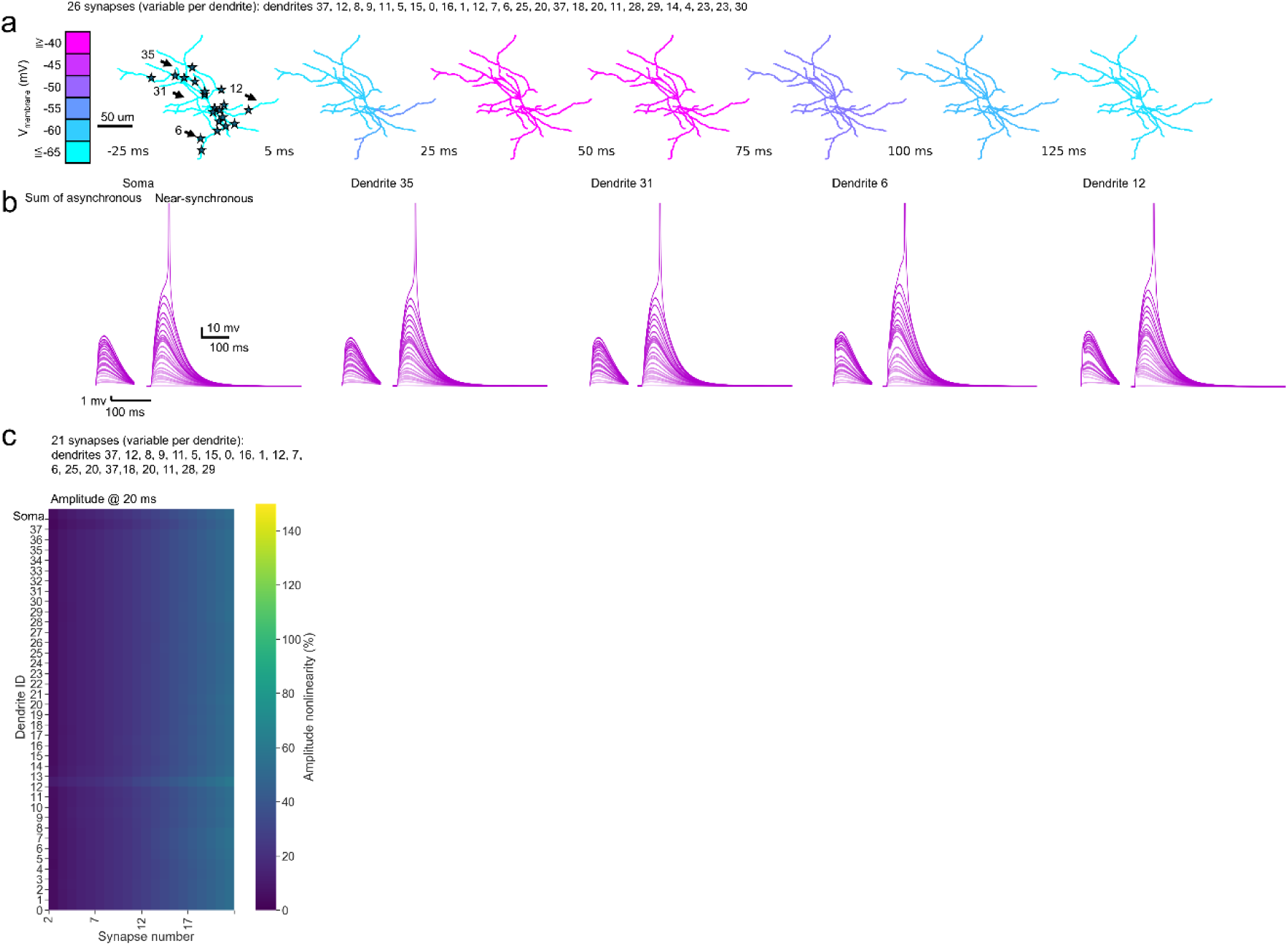
Spatially dispersed inputs can elicit dendritic supralinearities and somatic spikes. **(a)** Simulated dispersed synaptic input. 26 synapses (variable per dendrite) were activated on dendrites 37, 12, 8, 9, 11, 5, 15, 0, 16, 1, 12, 7, 6, 25, 20, 37, 18, 20, 11, 28, 29, 14, 4, 23, 23, 30. Voltage snapshots at 25 ms before stimulation and 5-125 ms after stimulation onset overlaid onto the neuronal architecture. Stimulation sites indicated by stars. Dendrites 35, 31, 6, 12 indicated with arrows have their uEPSPs displayed as traces below. **(b)** Sum of the asynchronous s and near-synchronous uEPSPs at the soma and dendrites 35, 31, 6, 12. **(c)** Heatmap of amplitude nonlinearity for every dendrite and the soma. Corresponds to (a, b). Maximum amplitude was measured.

## Materials and methods

The *ex vivo* voltage and calcium fluorescence data were obtained using methods described in Griesius et al. (22). The original data on neurogliaform cell LTP and associated methods were reported in Mercier et al. (30).

### Reconstruction

2-photon image stacks of neurogliaform interneurons, previously acquired and published (22), were used to generate morphological reconstructions for this study. Images were processed using Fiji (ImageJ 2.9.0), and neuronal morphologies were reconstructed using the Simple Neurite Tracer plugin (71). Reconstructed neurons were discretized into compartments representing the soma and dendrites. A total of three neurogliaform cell morphologies were reconstructed and used in subsequent simulations. Axonal arbors were not included. The same experimental dataset (22), including electrophysiological recordings and calcium imaging, was used to constrain and tune passive and active model parameters in the present study.

### Computational model

All simulations were performed using NEURON 7.7.2 software (44) and Python 3.7.4. The computational modelling code is available at the University College London repository at https://doi.org/10.5522/04/30104491.v2 under the CC BY-NC-ND 4.0 licence.

The morphology was adjusted so that the number of segments per dendrite was constrained to odd numbers and set according to the d-lambda rule (44). Passive parameters, including compartment diameter, resting membrane voltage (V_rest_), membrane resistance (R_membrane_), axial resistance (R_axial_), and capacitance, were set to approximate previously recorded experimental data for neurogliaform interneurons (Table 2) (30, 42). The morphological model was populated with dispersed active conductance mechanisms, including A-type and inward-rectifier potassium channels, voltage-gated sodium channels (VGSCs), L-type voltage-gated calcium channels (VGCC’s), and a leak conductance (I_leak_) (Table 3). VGSCs were inserted only into the soma to enable the model to generate action potentials, seeing as there was no evidence to suggest sodium-channel contribution to dendritic supralinearities (22). High-threshold L-type VGCCs were inserted into the dendrites, as this channel type was found to contribute to dendritic supralinearities in this cell type and generally (22, 50). A-type and delayed-rectifier potassium channels were inserted into the dendrites and soma as the likely candidate channels to counteract depolarization. Conductance densities, leak, and specific membrane capacitance were adjusted to match experimental data using key readouts like threshold, input resistance, and rheobase. The comparison between the intrinsic properties measured experimentally *ex vivo* and the outputs of the *in silico* computational model are summarised in Table 1.

**Table 2.** Initial model parameters.

| Parameter | Value |
| --- | --- |
| Dendrite diameter ($\mu\text{m}$) | 0.59-1.84 $\mu\text{m}$ |
| Soma diameter ($\mu\text{m}$) | 10 |
| $R_{\text{membrane}}$ kOhm $\text{cm}^2$ | 13.5 |
| $R_{\text{axial}}$ Ohm cm | 220 |
| $V_{\text{rest}}$ (mV) | -65 |
| Specific membrane capacitance<br>($\mu\text{F}/\text{cm}^2$) | 1.35 |
| Temperature ($^{\circ}\text{C}$) | 35 |
| Equilibrium potential (K) (mV) | -90 |
| Equilibrium potential (Na) (mV) | 55 |
| Equilibrium potential ($\text{Ca}^{2+}$) (mV) | 140 |

$R_{\text{membrane}}$ Membrane resistivity
$R_{\text{axial}}$ Axial resistivity
$V_{\text{rest}}$ Resting membrane voltage

**Table 3.** Peak conductance densities.

| Channel | Peak conductance (S/cm<sup>2</sup>) |  |
| --- | --- | --- |
|  | Soma | Dendrite |
| **Ka** | 0.016 | 0.008 |
| **Kdr** | 0.032 | 0.002 |
| **VGSC** | 0.425 | 0.001 |
| **VGCC (L-type)** | 0 | $1.15 \times 10^{-4}$ |
| **Passive conductance** | $7.40 \times 10^{-5}$ | $7.40 \times 10^{-5}$ |

Ka A-type potassium channel
Kdr delayed-rectifier potassium channel
VGCC voltage-gated calcium channel
VGSC voltage-gated sodium channel

AMPAR and NMDAR conductance-based point process mechanisms (45, 48) were inserted together to simulate glutamatergic synapses being activated (Table 4). The weights for both mechanisms were tuned to produce uEPSPs at the soma similar to those recorded experimentally, ∼1-2 mV in amplitude (22). The NMDAR weight was up to 4.5 times greater than the AMPAR weight to capture the large NMDAR-mediated currents in neurogliaform cells (30, 42). The synaptic weights and receptor rise and decay kinetics are similar to those in other studies investigating NMDAR-dependent dendritic supralinearities (14, 46, 47, 49, 50). Synaptic weights were varied randomly by a scaling factor of 0.2-1.8 to approximate the uEPSP variability observed experimentally during uncaging (22, 47). Modelled glutamatergic synapses were always made to include both AMPARs and NMDARs. Where multiple synapses were placed onto individual dendrites, synapses were always spread along the dendrites to approximate *ex vivo* uncaging input location arrangements; synapses were never superimposed.

**Table 4.** Synapse parameters.

| Parameter | Value |
| --- | --- |
| AMPA peak conductance ($\mu\text{S}$) | $2.5 \cdot 10^{-5}$ to $1.0 \cdot 10^{-4}$ |
| AMPA rise time constant (ms) | 0.5 |
| AMPA decay time constant (ms) | 1.5 |
| NMDAR peak conductance ($\mu\text{S}$) | $1.69 \cdot 10^{-5}$ to $6.75 \cdot 10^{-4}$ |
| NMDAR rise time constant (ms) | 4 |
| NMDAR decay time constant (ms) | 42 |

AMPAR AMPA receptor
NMDAR NMDA receptor

Total calcium current per compartment was calculated as the sum of up to two currents, depending on the compartment and on the where glutamatergic synapses were places: the VGCC current and a proportion of the NMDAR current. The dendritic NMDAR current at the sites of stimulation was captured and multiplied by the permeability of the NMDAR to calcium, estimated to be ∼7% (56–58). Total calcium current per compartment was calculated using the following equation:

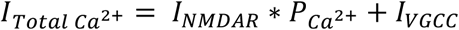

Where *I*_*NMDAR*_ is the NMDAR current, *P*_*ca*2+_ is the permeability of the NMDAR to calcium, and *I*_*VGCC*_ is the VGCC current.

In the simulations on clustered versus dispersed theta-burst like synaptic input, the focus was specifically on calcium influx via the NMDARs:

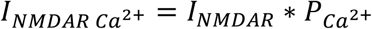

Where *I*_*NMDAR*_ is the NMDAR current, *P*_*ca*2+_ is the permeability of the NMDAR to calcium.

The different driving force for calcium than for monovalent cations was ignored for simplicity. Intracellular calcium dynamics arising such as calcium diffusion, endogenous buffering (59–63), and calcium-induced calcium release were not modelled here to avoid incorporating untested and unconstrained parameters for this cell type. The focus of calcium investigation was on calcium influx specifically in response to voltage change due to supralinear synaptic input summation and backpropagating action potentials.

The location of glutamatergic synapses, the timing of synaptic input, and the presence of somatic action potentials were varied across the simulations. The exact settings are described with the results for the respective experiments and are described in the computational modelling code. Randomization was performed using seeds to ensure reproducibility. Simulations used NEURON’s variable time-step solver (CVODE; cvode = 1). Absolute and relative error tolerances were set to *atol* = 0.001 and *rtol* = 0 (default settings). Recorded variables were sampled at 0.025 ms intervals; this sampling interval did not constrain the adaptive integration time step.

### Animals and husbandry

Adult male and female mice of varying ages were throughout the study (ages 1–3 months). Transgenic mouse lines were maintained as heterozygotes. *Ndnf^cre/cre^* or *Ndnf^cre/+^* mice (The Jackson Laboratory B6.Cg-*Ndnf^tm1.1(folA/cre)Hze^*/J; Stock No: 028536; Bar Harbor, ME, USA), bred on a C57BL/6 background were used to target neurogliaform interneurons (72). Animals were group-housed under a normal 12 hr light/dark cycle. Cages were enriched with paper bedding, cardboard tube, plastic tube, and wooden chewing blocks. Mice had had unlimited access to standard laboratory chow and water. Research was conducted in accordance with the Animal (Scientific Procedures) Act 1986 Amendment Regulations (SI 2012/3039) under PPL: PP3944290.

### Surgery for viral injection

Mice of at least 6 weeks of age were anaesthetized with isoflurane and placed into a stereotaxic frame, onto a heating pad to maintain body temperature. Mice were given Metacam (0.1 mg/kg) and buprenorphine (0.02 mg/kg) subcutaneously. Bilateral craniotomies were performed, positioned 3.1 mm caudal and ±3.12 mm lateral of Bregma. Virus (AAV2/9-mDlx-FLEX-mCherry; titre >1012 viral genomes/ml; VectorBuilder [Chicago, IL, USA]) was injected into the ventral CA1 region of both hippocampi using a Hamilton syringe 2.5 mm deep from the pia. 150 nL of virus was injected at each site at a rate of 100 nL/min. The needle was left in place for 5 min following injections before withdrawal. Mice were given 0.5 mL saline subcutaneously post-operatively to help with recovery and were monitored for 5 days following the procedure. Mice were sacrificed for experiments after a minimum of 3 weeks post-surgery.

### Brain slice preparation

Mice were sacrificed, the brains removed, and hippocampi dissected and sliced in ice-cold sucrose-based solution containing (in mM): 205 Sucrose, 10 Glucose, 26 NaHCO3, 2.5 KCl, 1.25 NaH2PO4, 0.5 CaCl2, 5 MgSO4, saturated with 95% O2 and 5% CO2. Transverse 400 µm hippocampal slices were cut using a Leica VT1200S vibrating microtome. Slices were incubated in artificial cerebrospinal fluid (aCSF) at 35 °C for 30 min and then at room temperature for 30 min. aCSF contained (in mM): mM: 124 NaCl, 3 KCl, 24 NaHCO3, 1.25 NaH2PO4 10 Glucose, 2.5 CaCl2, 1.3 MgSO4, saturated with 95% O2 and 5% CO2. The CA3 area was removed prior to slice transfer into the recording chamber to prevent recurrent activity.

### Electrophysiology

The recording chamber was perfused with oxygenated aCSF maintained at 32 °C. Slices were first visualized using Dodt illumination on an Olympus FV1000 BX61 microscope. Fluorescent cells were identified in the CA1 stratum lacunosum moleculare (neurogliaform cells) using Xcite epifluorescence. Borosilicate glass pipettes (pipette resistance of 4–5 MΩ) were pulled using a horizontal P2000 (Sutter Instruments) puller. The internal solution contained (in mM) 120 KMeSO3, 8 NaCl, 10 HEPES, 4 Mg-ATP, 0.3 Na-GTP, 10 KCl,∼295 mOsm, 7.4 pH. EGTA was not added to avoid introducing excess calcium buffering capacity. Internal solution aliquots were filtered on the day of experiment and small volumes of Alexa-Fluor-594 (final concentration 50 µM) and Fluo-4 (final concentration 400 µM) solution aliquots were added. Recordings were obtained using a Multiclamp 700B (Molecular Devices, USA) amplifier filtered at 10 kHz and digitized at 20 kHz (National Instruments PCI-6221, USA). WinWCP 5.5.6 (John Dempster, University of Strathclyde) software was used for data acquisition. After whole-cell break-in in V-clamp, the cells were switched to I=0 mode briefly to measure the resting membrane potential, before switching to I-clamp mode for experiments. All experiments were performed in I-clamp mode, with current continuously injected to maintain cell membrane between at ∼−65 mV (0 to –50 pA). All data are presented without adjustment for the junction potential (∼−15 mV). Recordings were discarded if they had an access resistance >25 MΩ and if access or input resistance changed by >20% over the course of the experiment. All experiments were conducted in the presence of picrotoxin (50 µM) and CGP55845 (1 µM) to block GABAergic transmission.

### Two-photon imaging

Two-photon imaging was performed using a Ti-sapphire lasers tuned to 810 nm (Mai-Tai, Spectra Physics). Cells were discarded upon observation of photodamage or upon change in resting membrane potential of >10 mV. All z-stacks used for the analysis of morphology and example figures were captured at the end of experiments to maximise the amount of time for experiments before cell health deterioration. Photomultiplier tube filters used: 515–560 nm (Fluo-4); 590–650 nm (Alexa-594).

### Quantification and statistical analyses of ex vivo data

Amplitude nonlinearity was regarded as the primary experimental outcome. For the purposes of quantifying nonlinearity, response amplitude was measured 20-50 ms after synaptic stimulation onset for most simulations, depending on experiment.

Response nonlinearity was quantified using the following equation(23, 73):

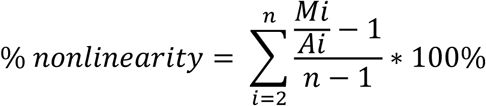

Where *Mi* is the amplitude of the *i*th measured uEPSP (composed of *i* individual stimulation spots), *Ai* is the amplitude of the *i*th constructed summed uEPSP, and n is the total number of stimulation locations.

Calcium signals were quantified using the following equation:

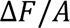

Where F is the fluorescence of the calcium indicator Fluo-4 and A is the fluorescence of the inert dye Alexa-594.

Voltage and calcium traces were filtered using a Savitzky-Golay filter. Uncaging light artefacts were removed from linescans. Due to the substantial variability in responses across dendrites, dendrites were used as the experimental unit. The numbers of dendrites, cells, and animals are reported in figure legends. α = 0.05 was applied for all statistical tests. The t and p values are presented in the figure legends and discussed in the text, as appropriate. Estimation statistics (74) and the generation of slopegraphs and Gardner-Altman estimation plots were performed using code provided by Ho et al., 2019 (75). The slopegraph indicates the difference in nonlinearity between predicted and recorded calcium fluorescence. The adjacent paired and unpaired mean difference plots are the group summaries. 95% confidence intervals of the mean difference were calculated by bootstrap resampling with 5000 resamples. The confidence interval is bias-corrected and accelerated. P values are provided with estimation statistics for legacy purposes only. Data were processed using custom Python code (76, 77), Microsoft Excel, and WinWCP 5.5.6 (John Dempster, University of Strathclyde). Figures were created using custom Python code, Microsoft PowerPoint, Procreate, ImageJ.

## Acknowledgements

We are grateful to Jonathan Cornford, Vincent Magloire, Kaiyu Zheng and other members of the Department of Epilepsy for help and advice.

## Contribution

SG: Conceptualization, Data curation, Investigation, Methodology, Writing – original draft, Writing – review and editing

AR: Resources, Investigation, Methodology, Writing – review and editing

MM: Investigation, Data curation, Writing – review and editing

DMK: Conceptualization, Supervision, Funding acquisition, Writing – original draft, Writing – review and editing

## Data availability

The 2-photon calcium imaging and LTP data and the code used for computational modelling are available at https://doi.org/10.5522/04/30104491.v2 under the CC BY-NC-ND 4.0 licence.

